# Ablation of glypican-3 enhances radiosensitivity in liver cancer by prolonging G2/M arrest and activating the ATM/CHK2 pathway

**DOI:** 10.64898/2026.05.11.724294

**Authors:** Joon-Yong Chung, Hima Makala, Woonghee Lee, Olivia W. Lee, Simran Khurana, Jeong Won Kim, Julia Sheehan-Klenk, Divya Nambiar, Stanley Fayn, Ayla O. White, Eun Joo Chung, Nada Alani, Sabrina Ramelli, Stephen M. Hewitt, Travis H. Stracker, Deborah E. Citrin, Peter L. Choyke, Freddy E. Escorcia

## Abstract

Glypican-3 (GPC3) is an oncofetal protein widely being explored as a diagnostic and therapeutic target in hepatocellular carcinoma (HCC). Given that radiotherapy in the form of external beam and radioembolization are standard-of-care treatments for HCC, we aimed to determine whether there was any relationship between GPC3 and response to radiotherapy. Here, we demonstrate that GPC3 expression confers radioresistance in liver cancer through integrated *in vitro, in vivo*, and patient-level clinical analyses. Stable GPC3-knockout in liver cancer cell lines (HepG2, Hep3B, Huh7) and ectopic GPC3 expression in GPC3-negative liver cancer cells (SNU449), as well as in non-hepatic A431 cells, demonstrated that GPC3-mediated radioresistance is not restricted to hepatic lineage. Following irradiation, GPC3-deficient cells exhibited reduced proliferation, impaired clonogenic survival, persistent DNA damage, prolonged G2/M arrest, and increased apoptosis. Transcriptomic profiling demonstrated enrichment of cell-cycle and DNA damage response pathways in irradiated GPC3-deficient cells compared with GPC3-positive cells, and protein analyses confirmed sustained activation of the ATM/CHK2 axis. *In vivo*, GPC3 deletion markedly enhanced radiation-induced tumor growth delay in both HepG2 and A431 xenograft models. Consistent with these findings, high GPC3 expression was associated with inferior clinical outcomes in patients with HCC undergoing external-beam radiotherapy or radioembolization. Together, these findings identify GPC3 as a determinant of radioresistance in liver cancer and suggest its potential utility as a biomarker to guide radiotherapeutic strategies.

**Significance statement:** Radiotherapy is an important treatment option for HCC, but biomarkers that predict tumor response remain limited. GPC3 is highly expressed in most HCCs and is being investigated as an important biomarker for diagnosis and treatment of this disease, yet its relationship, if any, on radiosensitivity has not been previously reported. Here, we identify GPC3 as a modulator of radioresistance. GPC3 loss enhances radiosensitivity and is associated with persistent unresolved DNA damage, prolonged G2/M arrest, and sustained activation of the ATM/CHK2 pathway, resulting in delayed tumor growth after irradiation. In a clinical cohort of patients treated with radiotherapy, high GPC3 expression was associated with poorer overall survival. These findings suggest that GPC3 expressing tumors may necessitate either more dose-intense radiotherapy, radiobioligically ablative and/or combined with other modalities, or alternative therapeutic modalities to adequately treat HCC.

## Introduction

Liver cancer is one of the leading causes of cancer-related mortality globally, with hepatocellular carcinoma (HCC) accounting for 80-90% of primary liver cancers (1). HCC is frequently diagnosed at an advanced stage due to the lack of early symptoms and limited adoption of screening protocols in high-risk individuals. Although HCC treatment has improved over the last few decades, the overall survival rate remains poor, with a five-year survival rate of 22% in the US (2). Liver transplantation and surgical resection can be curative for eligible patients. Locoregional treatments include thermal ablation, transarterial embolization approaches (bland-, chemo-, or radio-embolization), and external beam radiotherapy (EBRT), usually in the form of stereotactic body radiotherapy (SBRT). Hepatic artery infusion chemotherapy (HAIC) is also used, however, is usually limited to specialty centers (3). Systemic therapy, usually an immunotherapy-based regimen is typically reserved for patients with advanced disease (4). Recent technological advances in EBRT have played a pivotal role in allowing for excellent local control, and comparative studies have shown similar if not better outcomes when compared to embolization or thermal approaches (5–8). Additionally, like with other locoregional therapies (7, 9), EBRT can be used and is being systematically explored in combination with systemic therapies to enhance the treatment response and outcomes for patients with advanced HCC (10–13).

Given the important role that radiation therapy plays in HCC, as well as other cancers, it is critical to understand the mechanisms by which it exerts its therapeutic effects. Canonically, DNA has been considered the target of interest for radiotherapy, taking advantage of differential radiosensitivity of tumors compared to normal tissues due to impaired DNA damage repair or increased doubling time. Ionizing radiation (IR) activates the DNA damage response (DDR), a collection of pathways that recognize DNA lesions and activate cellular processes aimed at protecting genome integrity, including cell cycle checkpoint responses, DNA repair pathways and tumor suppressive programs, including apoptosis and senescence. The status of the DDR impacts cellular signaling, cell death pathway, and the immune microenvironment in both the irradiated tumor and surrounding healthy tissue (14). One of the most lethal forms of radiation-induced DNA damage is DNA double-strand breaks (DSBs) (15). DSBs are detected by the MRE11-RAD50-NBS1 (MRN) complex that recruits and activates the Ataxia Telangiectasia Mutated (ATM) kinase, the central transducer of the DDR to DSBs. ATM phosphorylates hundreds of downstream targets, including histone H2A family member X (H2AX), the checkpoint kinase CHK2, p53, breast cancer type 1 susceptibility protein (BRCA1) and radiation sensitive protein 51 (RAD51) (16–18). Due to their proximity to DSBs, phosphorylated histone H2AX (γH2AX) foci are widely used as a marker for monitoring radiation-induced DSB repair through immunofluorescent staining (19).

Cells employ two main pathways to repair DSBs: non-homologous end-joining (NHEJ) and homologous recombination (HR). NHEJ is the predominant DSB repair mechanism in mammalian cells, particularly during the G0/G1 phases of the cell cycle (20). NHEJ directly ligates the broken DNA ends without the need for a homologous template. In contrast, HR utilizes an undamaged sister chromatid as a template for accurate DSB repair, restricting its use to the S and G2 phases (21). The response to radiation therapy is influenced by a complex interplay of factors, including the tissue of tumor origin, tumor microenvironment, cell cycle status, DNA repair capacity, and inherent resistance mechanisms (22, 23).

Resistance to IR remains a significant obstacle for effective cancer treatment and specific markers that can reliably predict the efficacy of radiotherapy or recurrence after radiation therapy remain limited. This unmet need is especially relevant in HCC where there are several locoregional therapy options. Having a biomarker that could identify patients with tumors that would more likely benefit from one treatment versus another would be very valuable.

Glypican-3 (GPC3), a cell surface-associated heparan sulfate proteoglycan and oncofetal protein, plays a significant role in regulating various cellular processes, including growth, migration, and differentiation (24, 25). GPC3 is being explored as a diagnostic and therapeutic target for HCC because it is overexpressed in ∼75% of cases, where it is thought to contribute to tumor progression by influencing several signaling pathways (26). One of the key mechanisms by which GPC3 promotes HCC is by activating the canonical WNT signaling pathway (27, 28). This activation occurs through direct interaction between GPC3 and the Frizzled receptor, promoting tumor progression (27, 29). In addition to WNT signaling, GPC3 has been shown to modulate several other crucial pathways, including Hippo, YAP, AKT/PI3K, NF-κB, and MAPK/ERK (24, 25, 30–32). These pathways are closely tied to the cellular response to IR, and alterations in these pathways may affect radiosensitivity in HCC. Recent proteomics studies have also identified DNA damage-related proteins as potential GPC3 interactors (33), further suggesting that GPC3 could influence radioresistance. However, the mechanisms by which GPC3 contributes to radioresistance remain unclear.

Here, we tested the hypothesis that GPC3 expression confers resistance to IR. To address this, we engineered GPC3 knock-out and knock-in liver and non-liver cancer cell lines to systematically evaluate proliferation, clonogenic survival, DNA damage repair, and tumor growth in murine xenograft models. Furthermore, we investigated the clinical relevance of these findings by analyzing overall survival in a cohort of HCC patients treated with radiotherapy, stratified by GPC3 expression. Our results demonstrate that GPC3 expression is associated with increased radioresistance and accelerated DNA damage resolution. Notably, patients with GPC3^+^ HCC exhibited significantly poorer overall survival following radiotherapy compared to those with GPC3- tumors. Collectively, these data suggest that GPC3 serves as a predictive biomarker for radiotherapy efficacy in HCC and identifies a potential target for radiosensitization.

## Results

### GPC3 promotes tumor cell growth rates after irradiation

To investigate the impact of GPC3 on radioresponse in liver cancer cells, we genetically modified the expression levels of GPC3. HepG2 and Hep3B GPC3 knockout cell lines (GPC3^-^ HepG2 and Hep3B) were previously established using the CRISPR/Cas9 system (25). Additionally, an isogenic GPC3-overexpressing cell line derived from SNU449 (SNU449/GPC3) was created using a lentiviral expression system. Four GPC3^+^ (HepG2, Hep3B, SNU449/GPC3, and A431/GPC3) and four GPC3^-^ (GPC3^-^ HepG2 and Hep3B, SNU449, and A431) cell lines were assessed. Expression of *GPC3* mRNA and protein were confirmed in SNU449/GPC3, HepG2, and A431/GPC3 cell lines but were absent in SNU449, SNU449/Vector, GPC3^-^ HepG2, and A431 cell lines by qPCR and western blot assays (Fig. 1 *A* and *B*). Cell surface GPC3 expression was further examined in SNU449/GPC3, HepG2, and A431/GPC3 cell lines through flow cytometric analysis (Fig. 1*C*). GPC3 overexpression was also validated by immunofluorescence staining and clear GPC3 expression was evident in SNU449/GPC3 and A431/GPC3 but not in SNU449 and A431 cells (Fig. 1*D* and Supplementary Fig. S1).

**Fig. 1.**
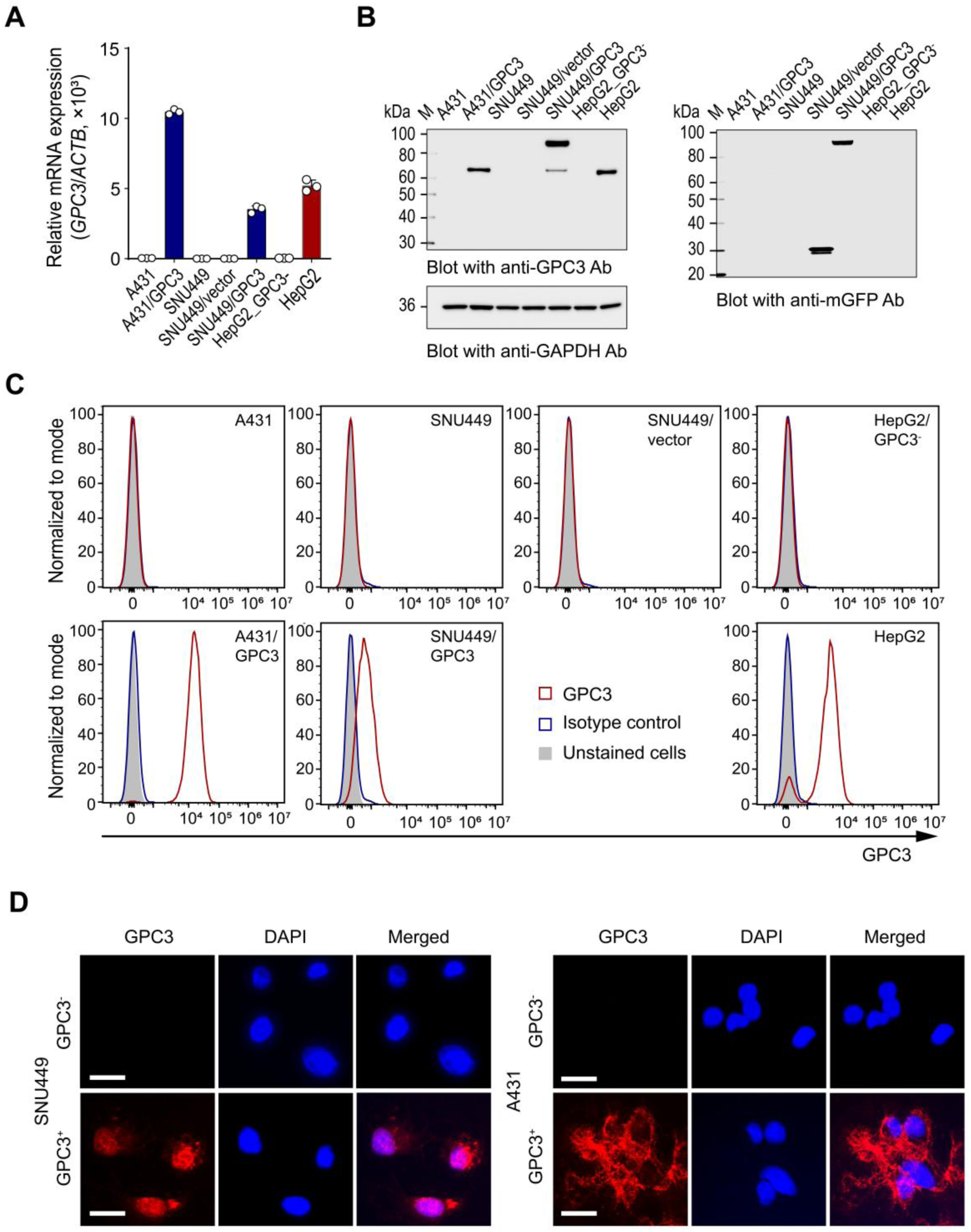
Ectopic GPC3 expression was successfully established and validated in isogenic SNU449 cells. (*A*) SNU449 cells were transduced with hGPC3-mGFP and clonally isolated, and GPC3 mRNA expression was quantified by qPCR. A431/GPC3 and HepG2 cells served as positive controls; A431 and GPC3⁻ HepG2 cells served as negative controls. (*B*) GPC3 protein expression was assessed by Western blotting (GAPDH loading control). (*C*) Cell-surface GPC3 was detected in A431/GPC3, SNU449/GPC3, and HepG2 cells, but not in GPC3⁻ SNU449, SNU449/vector, A431, or HepG2 cells. (*D*) Immunofluorescence staining confirmed GPC3 expression in SNU449/GPC3 and A431/GPC3 cells, but not in parental controls. Scale bars: 50 *μ*m.

In our prior study, we showed that GPC3 was required for increased tumorigenicity of liver cancer (25). To investigate the role of GPC3 on cell proliferation after IR, cell growth was recorded using an IncuCyte imaging system, which allowed for real-time monitoring of cells over a 9-day period following IR with 6 Gy. GPC3^-^ liver cancer cells displayed significantly reduced proliferation compared to their GPC3^+^ parental counterparts. By day 9, the proliferation rates of GPC3^-^ HepG2 and Hep3B were reduced by 70.7% and 33.4% relative to their parental cells, respectively (Fig. 2 *A* and *B*, and Supplementary Fig. S2). Similarly, GPC3^-^ Huh7 cells showed a 23.6% decrease in proliferation at day 6 post IR compared to GPC3^+^ Huh7 cells (Supplementary Fig. S3*A*). In contrast, GPC3^+^ SNU449 and A431 cells exhibited 2.0-fold and 9.5-fold increases in proliferation rates compared to their GPC3-deficient parental cells, respectively (Fig. 2 *C* and *D*, and Supplementary Fig. S2). These findings indicate that GPC3 expression level is positively correlated with tumor cell growth following IR.

**Fig. 2.**
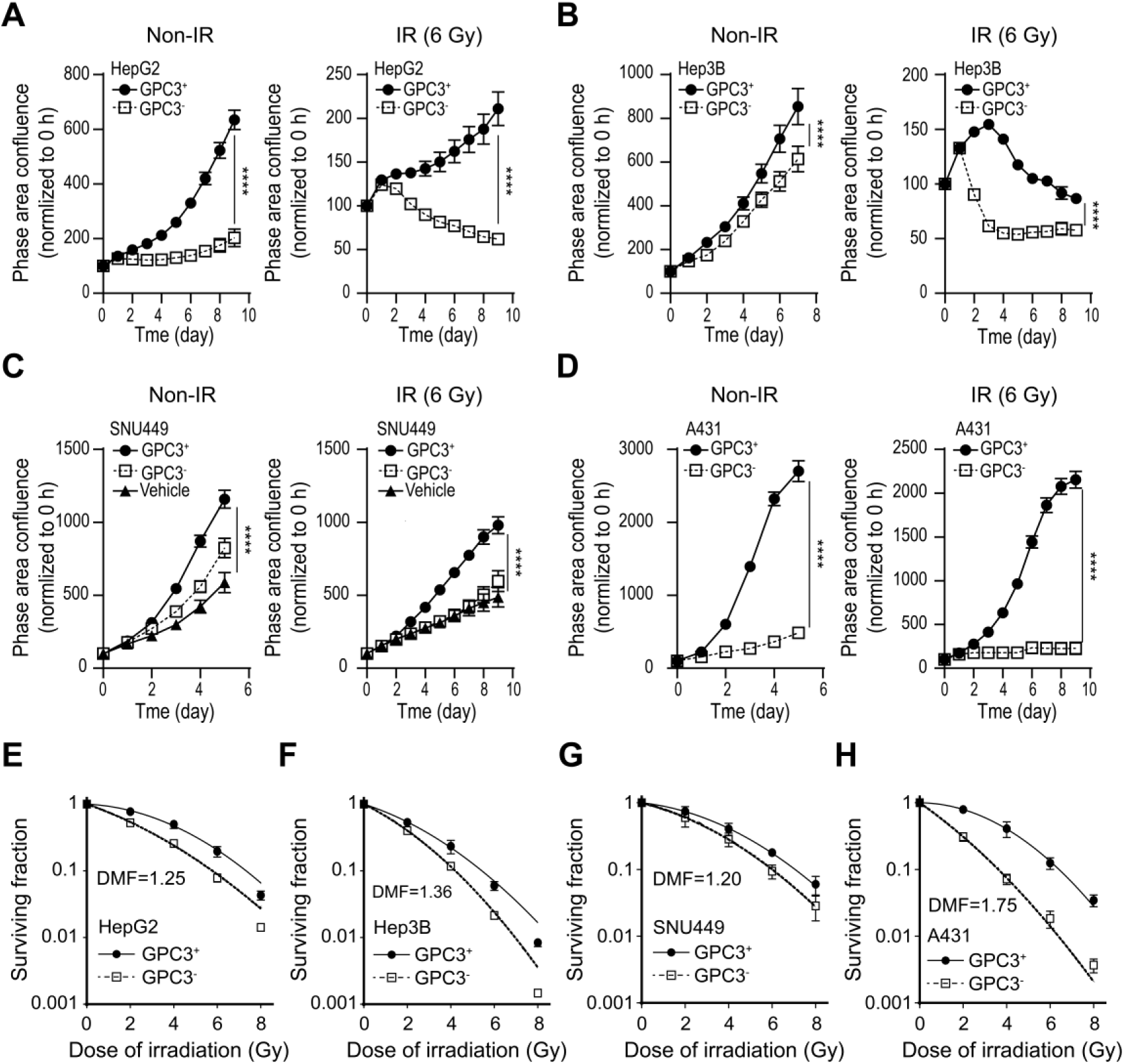
GPC3 expression regulates tumor cell proliferation and radiosensitivity. Proliferation of paired GPC3-positive (GPC3⁺) and GPC3-deficient (GPC3⁻) HepG2 (*A*), Hep3B (*B*), SNU449 and SNU449/V (empty vector control) (*C*), and A431 (*D*) cells was monitored using the IncuCyte® live-cell analysis system under non-irradiated (Non-IR) conditions and following 6 Gy irradiation (IR). Phase-area confluence was normalized to 0 h, and differences in proliferation kinetics were analyzed by two-way ANOVA, with significance assessed at the final time point (****, *P*<0.0001). Radiosensitivity was evaluated by clonogenic survival assays in the corresponding GPC3⁺ and GPC3⁻ HepG2 (*E*), Hep3B (*F*), SNU449 (*G*), and A431 (*H*) cells exposed to graded doses of γ-irradiation. Colonies were quantified 10–14 days later to generate survival curves. GPC3 loss increased radiosensitivity, yielding dose-modifying factors (DMFs) of 1.25 (HepG2), 1.36 (Hep3B), 1.20 (SNU449), and 1.75 (A431). Data represent mean surviving fraction ± SD from more than three independent experiments.

### GPC3 enhances clonogenic survival post IR in cancer cells

To explore the role of GPC3 on radiation-modulated clonogenicity in carcinoma cell, clonogenic survival assays were performed. In HepG2 and Hep3B liver cancer cell lines, GPC3 deficiency significantly reduced clonogenic survival following IR, with dose modifying factor (DMF) values of 1.25 and 1.36, respectively (Fig. 2 *E* and *F*). Conversely, GPC3 expression conferred radioresistance in GPC3^+^ SNU449 and GPC3^+^ A431 cell lines, with DMF values of 1.20 and 1.75, respectively (Fig. 2 *G* and *H*). Huh7 cells, which exhibit low endogenous GPC3 expression, showed only a marginal increase in survival post-irradiation (DMF = 1.16; Supplementary Fig. S3*B*), and were thus excluded from subsequent experiments. Overall, these findings highlight a dose-response relationship between GPC3 expression level and radioresistance in both liver cancers and epidermoid carcinoma cells by enhancing clonogenic potential.

### Radiation-induced DNA damage is prolonged in GPC3 deficient cancer cells

DNA repair plays a critical role in the effectiveness of radiation therapy by influencing the survival of both cancer and normal cells. To examine DSB induction and repair, we assessed the formation of γH2AX nuclear foci, a widely recognized biomarker for detecting and quantifying DSBs (19). Cells were seeded and irradiated with a dose of 6 Gy and γH2AX foci were quantified at 1, 6, 24, and 48 h post-IR. IR led to a significant increase in γH2AX foci in both GPC3^+^ and GPC3^-^ cell lines at 1 h after IR. However, the rate of foci resolution varied depending on GPC3 expression levels. In GPC3^-^ cells, γH2AX foci remained significantly elevated compared to their parental cells, with this effect persisting from 6 to 48 h post-IR (Fig. 3 *A* and *B*). In contrast, GPC3^+^ cells exhibited a marked reduction in foci numbers relative to GPC3^-^ cells, such as SNU449 and A431 (Fig. 3 *C* and *D*).

**Fig. 3.**
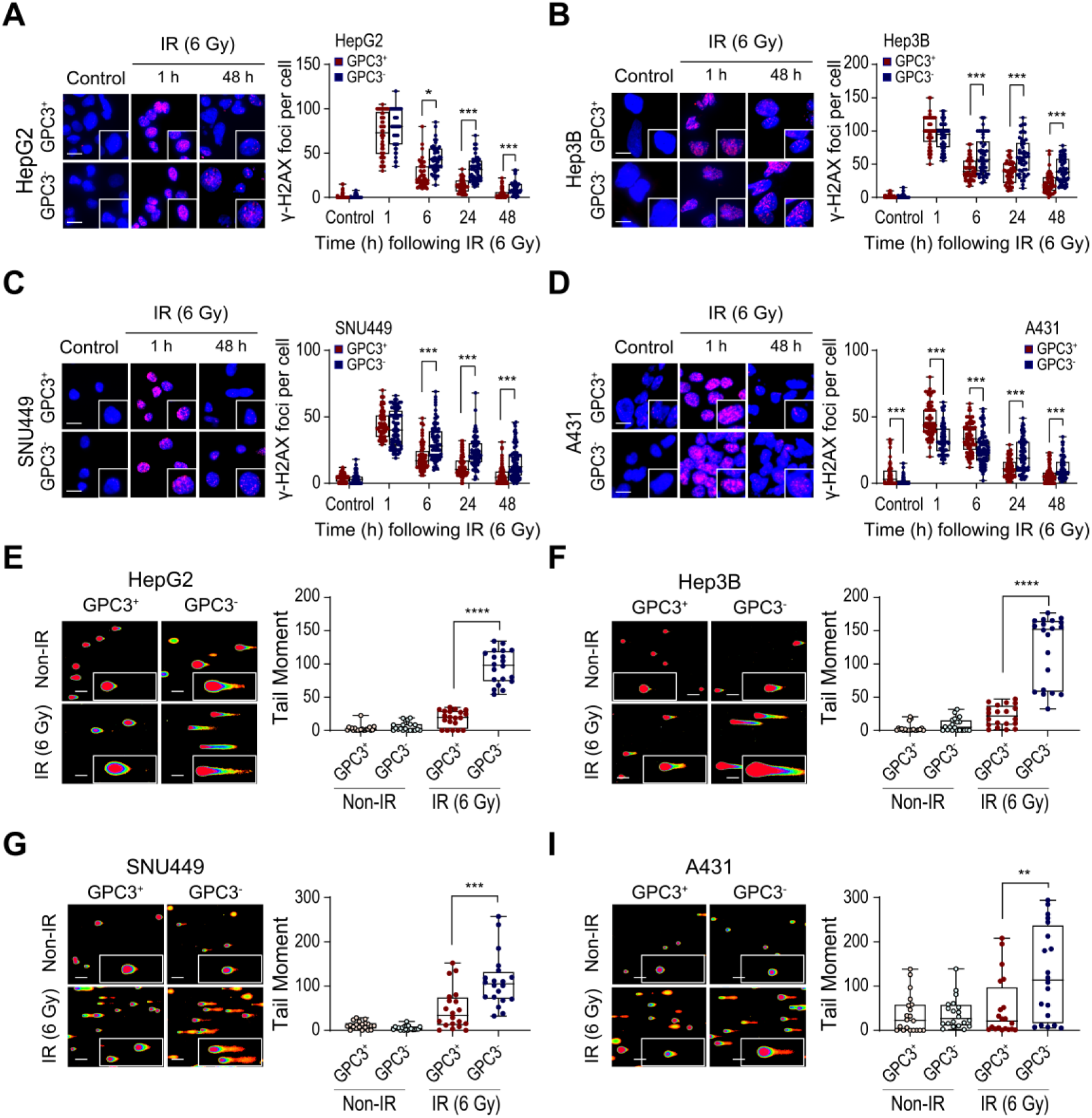
GPC3 deficiency increases IR-induced DNA double-strand breaks in tumor cells. DNA damage responses were assessed in paired GPC3-positive (GPC3⁺) and GPC3-deficient (GPC3⁻) HepG2 (*A*, *E*), Hep3B (*B*, *F*), SNU449 (*C*, *G*), and A431 (*D*, *I*) cells following 6 Gy irradiation (IR). γ-H2AX immunocytochemistry (*A*–*D*) was performed at the indicated time points to quantify DNA double-strand break formation and resolution. Representative images at control, 1 h, and 48 h post-IR are shown. γ-H2AX foci (magenta) were quantified from > 50 cells per condition; nuclei were counterstained with DAPI (blue). Scale bars: 25 *μ*m. Statistical comparisons of foci numbers were performed using a *t*-test (\**p*<0.05; \*\*\**p*<0.001). Comet assays (*E*–*I*) were performed 24 h after IR. Representative comet images (left) and quantified tail moments (right) are shown. Data are presented as mean ± SD, and significance was determined using a *t*-test (\*\**p*<0.01; \*\*\**p*<0.001; \*\*\*\**p*<0.0001). Scale bars: 50 *μ*m.

To further evaluate the genotoxic impact of GPC3 on DNA repair following IR, we utilized the alkaline comet assay. GPC3^-^ cells exhibited longer tail moments compared to their GPC3^+^ cells (Fig. 3 *E-I*). Overall, these data suggest that GPC3-deficient cells have delayed DNA repair rates reflected by significantly higher γH2AX foci retention and longer tail moments after IR compared to GPC3^+^ cells.

### GPC3 deficiency enhances late apoptosis and G2/M phase arrest in liver cancer cells post irradiation

Because radiation-induced cell cycle checkpoint arrest and apoptosis are influenced by the DNA repair process, we investigated the effect of GPC3 deficiency on cell cycle progression and apoptosis following IR (6 Gy) in liver cancer cells using flow cytometry. In GPC3^-^ HepG2 cells, the proportion of cells in the G2/M phase at 48 h after IR was comparable to that in GPC3^+^ cells (29.1% vs. 31.1%). However, at 72 h post IR, GPC3^-^HepG2 cells exhibited a significantly higher fraction of G2/M phase cells (34.9%) than GPC3+ cells (27.5%). In the absence of IR, GPC3^-^ HepG2 cells also displayed an increased G2/M fraction at 72 h (43.0%) relative to GPC3^+^ cells (30.2%) (Fig. 4*A*). Similarly, GPC3^-^Hep3B cells showed an increased proportion of cells in the G2/M phase following IR, with 74.1% and 48.0% at 48 and 72 h post-IR, respectively, compared with 57.4% and 33.4% in GPC3^+^ cells (Fig. 4*B*). We next assessed the impact of GPC3 deficiency on apoptosis in the presence or absence of IR. In HepG2 cells, GPC3 deficiency modestly increased late apoptosis and necrosis at 72 h post-IR, rising from 7.4% and 7.0% in GPC3^+^ HepG2 cells to 12.0% and 12.2% in GPC3^-^ HepG2 cells, respectively (Fig. 4*C*). No significant differences were observed at 24 or 48 h post-IR (Supplementary Fig. S4 *A* and *B*). Similarly, late apoptotic populations were higher in GPC3^-^ Hep3B cells under both non-irradiated (7.6% vs. 4.8%) and irradiated (17.4% vs. 14.4%) conditions at 72 h post-IR, compared with parental cells (Fig. 4*D*). Consistent with observations in HepG2 cells, no meaningful differences in early apoptosis, late apoptosis, or necrosis were detected at 24 or 48 h post-IR between GPC3^-^ and GPC3^+^ Hep3B cells (Supplementary Fig. S4 *C* and *D*). These findings suggest that GPC3 deficiency sensitizes cells to IR and alters DNA damage responses, leading to sustained G2/M cell cycle arrest and increased apoptotic cell death.

**Fig. 4.**
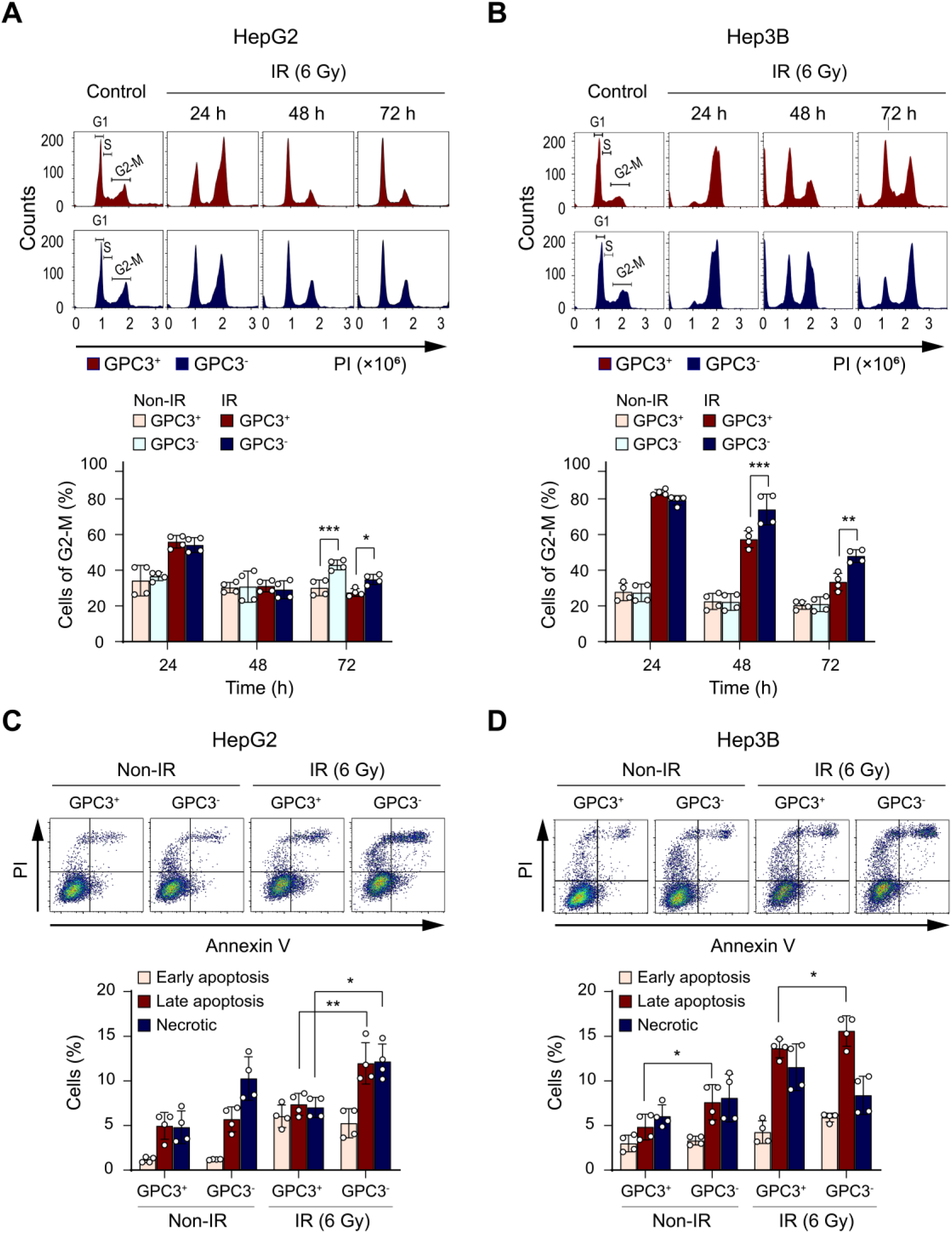
Irradiation induces distinct cell-cycle responses and cell-death in GPC3⁺ and GPC3⁻ liver cancer cells. (*A*, *B*) Cell-cycle profiles of irradiated GPC3⁺ and GPC3⁻ HepG2 and Hep3B cells were analyzed by PI staining at the indicated time points. Representative histograms and summary data (mean ± SD) are presented. Statistical significance was determined using a *t*-test (\**p*<0.05; \*\**p*<0.01; \*\*\**p*<0.001). (*C*, *D*) GPC3⁺ and GPC3⁻ HepG2 and Hep3B cells were exposed to 6 Gy irradiation (IR), and cell death was assessed by Annexin V/Propidium Iodide (PI) staining. Representative dot plots are shown, with quantification from three independent experiments (mean ± SD).

### GPC3 deletion promotes sustained DNA damage signaling and cell-cycle arrest following irradiation in liver cancer cells

To investigate the mechanism by which GPC3 influences DDR and tumor radiosensitivity, we performed RNA-seq to identify differentially expressed genes (DEGs) in HepG2 and Hep3B cells following IR. In GPC3-deficient HepG2 and Hep3B cells, we found 1371 and 3035 upregulated DEGs, and 455 and 1049 downregulated DEGs, compared with their parental GPC3-expression countparts, respectively (Supplementary Fig. S5 *A* and *B*). Ingenuity Pathway Analysis (IPA) identified significant changes in molecular and cellular functions, including cell cycle regulation, DNA replication, recombination, and repair, in GPC3^-^ cells post-IR (Fig. 5*A*). To further delineate the molecular pathways associated with GPC3 deficiency following IR, we performed gene set enrichment analysis (GSEA). We observed that pathways including cell localization, differentiation, and migration were all downregulated in GPC3^-^ HepG2 cells (Fig. 5*B*). Similarly, GPC3^-^ Hep3B cells exhibited downregulation of cell cycle-related pathway genes, further corroborating the role of GPC3 in cell cycle regulation after irradiation. Given the relatively limited sample size and the potential for statistical overcorrection in the differential expression analysis, we also examined the normalized raw counts of key genes associated with functions identified in IPA. Consistent with our pathway analyses, we observed differential expression of several genes involved in the cell cycle between GPC3-deficient and parental cell lines (Fig. 5*C*). Interestingly, HepG2 and Hep3B cells exhibited differences in the expression levels of specific genes involved in the top altered pathways, suggesting cell line-specific variations in the molecular response to GPC3 deficiency and IR.

**Fig. 5.**
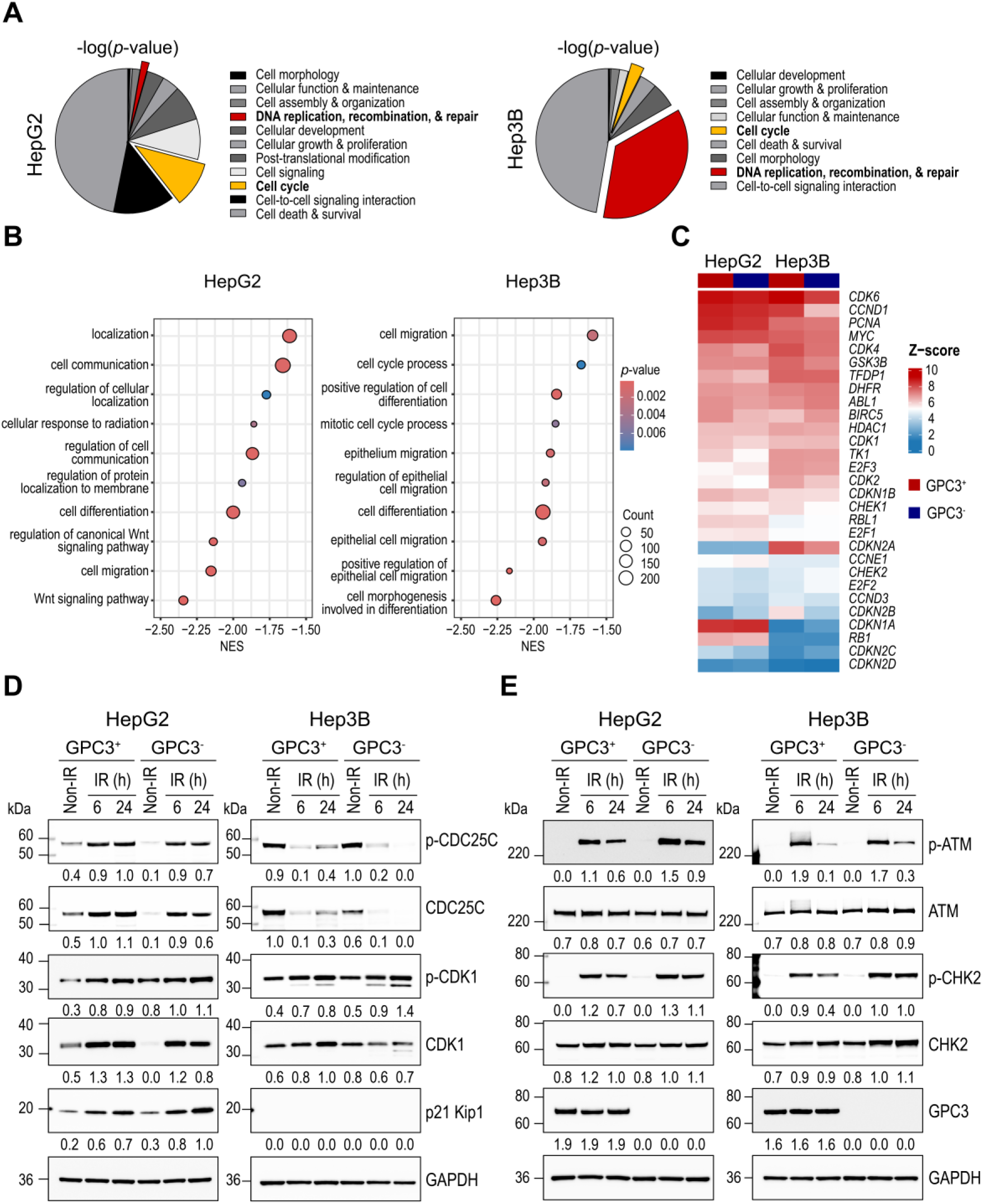
GPC3 deficiency alters transcriptional programs and signaling pathways in liver cancer cells following irradiation. (*A*) Ingenuity Pathway Analysis (IPA) identified molecular and cellular functions differentially regulated in GPC3⁻ versus GPC3⁺ liver cancer cells (all *p* < 2.09 × 10⁻⁵). (*B*) Dot plot of enriched KEGG pathways and Gene Ontology Biological Process (GOBP) terms in GPC3⁻ cell lines compared with parental controls, with normalized enrichment score (NES), gene count, and adjusted *p-*values indicated. (*C*) Heatmap showing differential expression of cell cycle–related genes in GPC3⁻ cells. (*D*, *E*) Total cell lysates collected at 6 and 24 h after irradiation (IR) were analyzed by Western blot for G2/M arrest– and cell cycle–related proteins (*D*) and ATM/CHK2 pathway components (*E*). Non-irradiated samples served as controls, and fold-change values are shown below each blot. GAPDH was used as a loading control.

### GPC3 deficiency induces sustained G2/M arrest through ATM/CHK2/CDC25C pathway activation in irradiated liver cancer cells

Based on the flow cytometry and RNA-Seq data analysis, which demonstrated significant alterations in DNA repair and cell cycle progression following IR, we analyzed the protein levels of key regulators of cell cycle progression in liver cancer cells. Among the cell cycle-related proteins, total CDC25C, and phosphorylated CDC25C (p-CDC25C) levels were significantly reduced 24 h post IR in GPC3^-^ liver cancer cells compared with GPC3+ cells. Conversely, total CDK1 levels were elevated, while phosphorylated CDK1 (p-CDK1) levels were decreased under the same conditions. In addition, the expression of p27 Kip1 (*CDKN1B*), a negative regulator of cell proliferation, was notably increased in GPC3^-^HepG2 cells, whereas it was undetectable in Hep3B cells (Fig. 5*D*). The increased phosphorylation of CDK1 at Y15 and the downregulation of CDC25C are hallmark indicators of G2/M phase arrest.

To further explore the mechanisms underlying GPC3^-^ induced G2/M phase arrest, we examined the ATM/CHK2 signaling pathway that is an upstream regulator of CDC25C. We found that phosphorylated S1981-ATM (p-ATM) and phosphorylated T68-CHK2 (p-CHK2) levels were markedly elevated following IR, reflecting their activation by DNA damage, followed by a gradual decrease over time (Fig. 5*E*). Importantly, the levels of p-ATM remained higher 24 h post-IR in GPC3^-^ liver cancer cells compared to the parental controls. These findings further support the observations that DNA repair is delayed in GPC3^-^ cells, leading to prolonged signaling through ATM/CHK2/CDC25C that control the G2 checkpoint and apoptotic responses.

### GPC3 knockout improves the efficacy of irradiation in xenograft tumor models

Given the observed increase in radiosensitivity of tumor cells *in vitro* following GPC3^-^, we extended our study to *in vivo* xenograft models using athymic nude mice. HepG2 and A431 tumor cells were selected for these xenograft models, as GPC3^-^ Hep3B and SNU449 cells failed to establish tumors. Three single-cell clones (H2K17, T2, and AL1 for HepG2-KO/Luc, GPC3^+^ A431 (G1)/Luc, and A431/Luc, respectively) were selected for further analyses (Supplementary Fig. S6).

GPC3^+^ and GPC3^-^ HepG2/Luc cells were injected subcutaneously implanted into the hind limbs of nude mice, using 5×10⁶ GPC3⁺ and 7×10⁶ GPC3^-^ cells per injection. Radiation therapy was administered once tumors reached approximately 150–200 mm³ in volume. A significant tumor growth delay was observed in GPC3^-^ tumors treated with radiation, while untreated GPC3⁺ and GPC3⁻ tumors exhibited comparable growth kinetics (Fig. 6 *A* and *B*, and Supplementary Fig. S7 *A*-*C*). By day 28 post IR, the average tumor volume in the GPC3^+^ HepG2/Luc group was 1590 ± 126 mm³, while the GPC3^-^HepG2/Luc group exhibited a significantly smaller average tumor volume of 337 ± 101 mm³, representing a 78.8% reduction compared to GPC3^+^ HepG2/Luc tumors (*p*<0.0001, Fig. 6*C* and Supplementary Fig. S7*C*). Notably, no significant differences in body weight were observed between the GPC3^+^ and GPC3^-^ groups throughout the experimental period (Supplementary Fig. S7*D*). Immunohistochemical analyses confirmed cell surface GPC3 expression was exclusively detected in GPC3^+^ cells. Additionally, the expression of Ki-67, a marker for cell proliferation, was markedly lower in GPC3^-^ tumors compared to GPC3^+^ tumors that were treated with radiation (Fig. 6*D*).

**Fig. 6.**
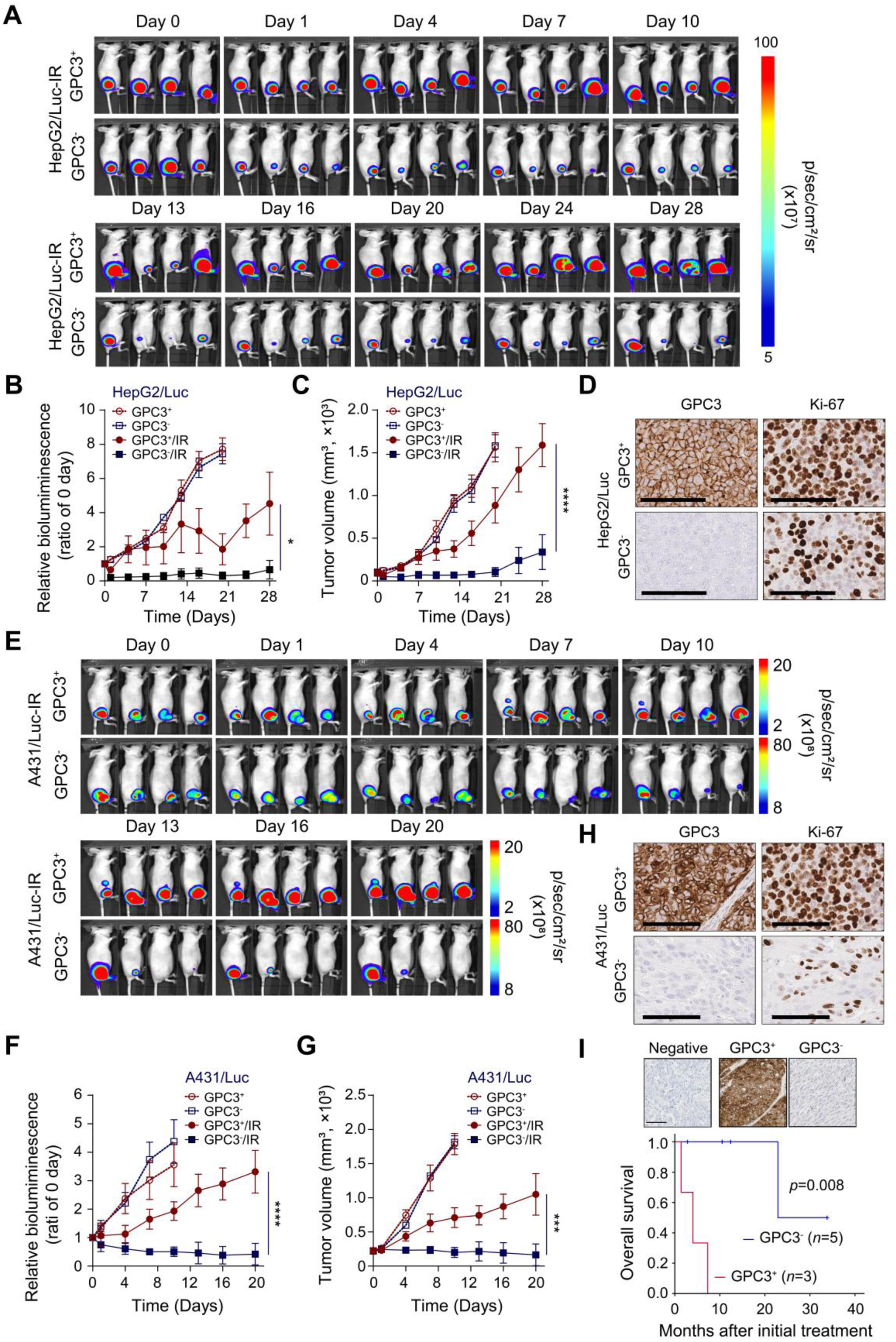
GPC3 deficiency enhances radiation response in xenograft models and correlates with clinical outcomes. (*A*–*D*) HepG2/Luc xenografts derived from GPC3⁺ or GPC3⁻ cells with or without 10 Gy irradiation were monitored by bioluminescence imaging (BLI) and tumor volume measurements. GPC3⁻ tumors showed enhanced growth delay following irradiation. Representative GPC3 and Ki-67 staining is shown (scale bar: 100 *μ*m). (*E*–*H*) Similar analyses in A431/Luc xenografts demonstrated greater radiation sensitivity in GPC3⁻ tumors. (*I*) In a clinical cohort, high tumor GPC3 expression was associated with poorer overall survival (*p* = 0.008). Statistical comparisons were performed by two-way ANOVA; significance was assessed at the final time point (\**p*<0.05; \*\*\**p*<0.001; \*\*\*\**p*<0.0001).

To further confirm the generalizability of the effect of GPC3 on radiosensitivity *in vivo*, similar experiments were conducted using GPC3^+^ and GPC3^-^ A431/Luc xenografts. Consistent with the findings in HepG2 models, GPC3^-^ tumor xenografts exhibited significantly delayed growth compared to GPC3^+^ xenografts (Fig. 6 *E* and *F*, and Supplementary, Fig. S7 *E-G*). By day 24 post IR, the average tumor volume of GPC3^+^ xenografts (G1) reached 1048 ± 101 mm³, while GPC3^-^ xenografts (A431) measured 164 ± 53 mm³, a reduction of 84.4% compared to GPC3^+^ tumors (*p*<0.001, Fig. 6*G* and Supplementary Fig. S7*G*). As in the HepG2 model, no significant body weight differences were observed between GPC3^+^ and GPC3^-^ groups (Supplementary Fig. S7*H*). Immunohistochemistry confirmed GPC3 expression exclusively in GPC3^+^ A431 tumors, and Ki-67 expression was significantly higher in GPC3^+^ tumors following IR (Fig. 6*H*). These findings collectively indicate that GPC3 deletion significantly enhances tumor growth delay in HepG2 and A431 xenografts following radiation therapy.

### GPC3^+^ HCC patients exhibit poorer overall survival rates following radiation therapy compared to GPC3^-^ HCC patients

Building on the finding that GPC3 deficiency enhances radiosensitivity in both *in vitro* and *in vivo* models, we investigated whether GPC3 expression influences the response to radiation therapy in HCC patients. While few patients with HCC undergo biopsies and radiotherapy has only recently become more commonplace for HCC treatment, we were able to obtain and analyze 8 cases of patients who underwent EBRT. Among the analyzed HCC specimens, 5 out of 8 cases (62.5%) exhibited low GPC3 expression (GPC3^-^, average histoscore 0.22), while 3 cases (37.5%) showed high GPC3 expression (GPC3^+^, average histoscore 45.8; Supplementary Table S1). Survival analysis using a Kaplan-Meier plot demonstrated that patients with GPC3^+^ tumors exhibited significantly worse overall survival (OS) compared with GPC3^-^ tumors (median OS 4.0 months vs. not reached, respectively; log-rank *p*=0.008; Fig. 6*I*). At last follow-up, all patients with GPC3^+^ tumors had died, whereas 80% of patients with GPC3^-^ tumors remained alive. These findings suggest that GPC3 expression is associated with poorer outcomes in patients with HCC who receive radiotherapy that is not radiobiologically ablative and GPC3 positivity may reflect more aggressive biology and early metastatic disease, perhaps necessitating comparatively earlier deployment of systemic therapy.

## Discussion

GPC3 is actively being studied as a biomarker to facilitate the diagnosis, prognosis, and therapeutic targeting of HCC. To date, however, its role in radiation response has been understudied, despite the routine use of radiotherapy (EBRT or radioembolization) in clinical practice (34, 35). In the current study, we demonstrate that GPC3 expression is associated with the radioresistance of liver cancer cells using *in vitro* and *in vivo* models. We also found that patients with GPC3^+^ HCC exhibited poorer survival outcomes compared to those with GPC3^-^ HCC after radiotherapy. Our results suggest that in patients with HCC who received radiotherapy, GPC3 predicts poorer outcomes, suggesting both that personalized approaches may be warranted and that targeting GPC3 could have potential therapeutic benefits. However, larger prospective studies are needed for further validation.

The DDR is a vital cellular mechanism responsible for detecting, signaling, and repairing damaged DNA. These pathways are activated when DNA damage is detected, triggering cell cycle arrest and the recruitment of specific repair machineries tailored to the nature of the lesion. Our findings indicate that GPC3 functions as a biological modulator of the DDR, particularly in the orchestration of DSB repair. While cells employ multiple pathways such as NHEJ or HR depending on the cell cycle phase, GPC3 expression appears to promote the efficiency of these processes, likely via the ATM/CHK2 axis. Consistently, GPC3 deficient cells exhibited significantly lower proliferation rates, reduced clonogenic survival and elevated apoptosis following IR. The accumulation of unrepaired DSBs and prolonged G2/M arrest, coupled with persistent ATM/CHK2 signaling in GPC3^-^ cells, collectively suggest that GPC3 loss compromises the kinetics and fidelity of DNA repair.

In response to DNA damage, activated CHK2 inhibits the activity of CDC25C, which subsequently suppresses downstream CDK1, (36), leading to G2/M phase arrest. In agreement with this mechanism, GPC3^-^ liver cancer cells demonstrated increased levels of inactive CDK1 (phosphorylated form), along with a marked reduction in total CDK1 (active form) expression. The levels of inactive or active CDC25 were notably lower in GPC3^-^ liver cancer cells compared to GPC3^+^ liver cancer cells. We also confirmed prolonged G2/M arrest in response to IR in GPC3^-^ liver cancer cells by flow cytometry. These findings suggest that the reduced levels of CDC25C in GPC3 deficient cells fail to adequately dephosphorylate CDK1, thereby contributing to the G2/M arrest prolongation observed in these cells. G2/M arrest can be regulated through p53-dependent (p53-p21-CDK1) and p53-independent (CHK1/2-CDC25C/CDK1) pathways (37). The upregulation of p21, in combination with p53, extends G2/M arrest following DNA damage (38). However, G2/M arrest can also occur in the absence of functional p53.

In this study, we examined the level of p21 expression in GPC3^+^ and GPC3^-^ HepG2 cells following IR. Our results showed a significant increase in p21 expression in GPC3^-^HepG2 cells compared to GPC3^+^ parental cells. Conversely, no detectable p21 expression was observed in Hep3B cells. This is consistent with a previous report that G2/M arrest induced by ATM-CHK2 does not impact p21 expression. Interestingly, Hep3B cells, which inherently lack p53 expression, without p21 expression exhibit a greater degree of G2/M arrest compared to HepG2 cells with high levels of p21. Thus, we conclude that G2/M arrest in GPC3^-^ liver cancer cells following IR occurs through both p53-dependent and p53-independent mechanisms, irrespective of p21 expression levels. Further studies are needed to explore the detailed molecular mechanisms through which GPC3 influences the DNA damage repair process, particularly in relation to p53 and p21 expression status.

In the current era of biomarker-driven cancer therapy, extensive efforts are being made to identify ones that can guide selection of appropriate therapies and predict therapeutic outcomes across various cancers, including HCC (39). These biomarkers possess significant potential for understanding tumor biology, which is crucial for effective clinical management. However, integrating biomarkers into radiotherapy guidance remains challenging due to limited data as well as the gap between basic research findings and clinical application. This challenge is particularly pronounced in HCC. In this study, tumor growth was significantly delayed in GPC3^-^ xenograft tumors with following IR. Additionally, patients with HCC harboring low levels of GPC3 showed increased overall survival compared to patients with high GPC3 expression. Although we measured GPC3 expression in tissue samples from patients, serum GPC3 levels are also being explored as a potential complementary biomarker to serum alpha-fetoprotein for diagnosing, surveilling, and monitoring treatment response of patients with HCC (40). However, further studies are needed to clarify the role of GPC3 as a predictive biomarker for radioresistance by examining its levels in both blood and tissue samples from a larger cohort of HCC patients, ideally in a prospective study.

Due to its high and specific expression in HCC, and promising findings from preclinical and clinical studies, GPC3 has emerged as an appealing target for both diagnostic and therapeutic strategies in this disease. Recent developments have introduced various GPC3-targeted therapies for managing HCC, including specific antibodies against GPC3 (41), antibody-drug conjugates (42), peptide and DNA-based vaccines (43, 44), and chimeric antigen receptor (CAR) T-cell (45, 46). Specifically, radiopharmaceutical diagnostics have been shown to successfully identify GPC3-positive lesions and monitor treatment responses (47–51), providing a non-invasive tool for patient stratification. In parallel, preclinical radiopharmaceutical therapies have demonstrated the ability to effectively inhibit tumor growth and enhance survival by selectively targeting GPC3-expressing cells. These encouraging outcomes have led to clinical trials investigating the safety, biodistribution, and therapeutic potential of these agents in patients with advanced HCC (52, 53). Despite these advantages, challenges persist, such as optimizing the choice of diagnostic and therapeutic radioisotopes, improving delivery efficiency, and mitigating potential off-target effects or radiation-induced toxicity.

In the present study, our research explores the potential role of GPC3 in radiotherapy. Our findings suggest that GPC3 expression may confer resistance to radiation. These insights could have implications for standard treatment practices. For instance, GPC3-positive tumors could benefit from radiobiologically ablative doses, whereas less aggressive dosing may suffice for GPC3-negative tumors. Additionally, one could envision that patients bearing GPC3-positive tumors may require a lower threshold to use systemic therapies given increased aggressiveness and higher metastatic potential. This approach would necessitate a shift in current practice, as routine biopsies are not typically performed in HCC due to the high diagnostic accuracy of non-invasive imaging techniques like multiphase computed tomography (CT) and magnetic resonance imaging (MRI). However, tissue biopsies could enable more precise characterization of disease at the patient level, facilitating better selection for localized or systemic therapies. Although the risk of tumor seeding from needle biopsies is a valid concern, a recent systematic review indicated that the reported risk is relatively low, 0.62 % (54).

Our study has several limitations. First, our experiments were restricted to *in vivo* models using hepatoblastoma and epidermoid carcinoma cells due to the limited availability of isogenic, paired HCC cells that could generate tumors. Second, the validation of the *in vitro* and *in vivo* findings was conducted in clinical samples from a single institution with a limited cohort size and was not designed as a prospective validation study. Therefore, we consider the present work as hypothesis generating, worthy of pursuing further study.

Here, we demonstrate, for the first time, that GPC3 expression is associated with promoting IR-induced DNA damage repair and radioresistance *in vitro* and *in vivo*. Furthermore, we confirm that, in a small retrospective clinical cohort of patients treated with either external radiotherapy or radioembolization, patients with GPC3-expressing HCC exhibit poorer overall survival compared to those with GPC3-negative HCC.

## Materials and Methods

### Cell lines & treatments

Human liver cancer cell lines (HepG2, Hep3B, & SNU449) and the cutaneous squamous cell carcinoma line (A431) were obtained from the American Type Culture Collection (ATCC, Manassas, VA, USA). Genetically modified liver cancer cell lines with knock-out or overexpression GPC3 were generated in-house. The Huh7 and GPC3^+^ A431 cell line (aka, G1) was provided by Dr. Mitchell Ho (Bethesda, USA). The HepG2-Luciferase (Luc) cell line was purchased from Revvity (Waltham, MA, USA), while we generated GPC3^-^HepG2/Luc, GPC3^+^ and GPC3^-^ A431/Luc cell lines for this study. Cells were cultured in DMEM (Thermo Fisher Scientific, Carlsbad, CA, USA), RPMI 1640 medium (Thermo Fisher Scientific), or EMEM (ATCC), each supplemented with 10% fetal bovine serum (FBS; Thermo Fisher Scientific), in a humidified atmosphere of 5% CO_2_ at 37℃. Cells grown to 70–80% confluence in 10 cm dishes were irradiated using a 320 kV X-ray source (Precision X-Ray Inc., North Branford, CT, USA) at a dose rate of 2.3 Gy/min. Mycoplasma contamination was ruled out in all cell lines using the MycoAlert Mycoplasma Detection Kit (Lonza, Basel, Switzerland).

### Patient specimens

This study retrospectively analyzed tissue samples from eight individuals diagnosed with HCC who underwent radiotherapy between September 2013 and December 2020 at Kangnam Sacred Heart Hospital. Patients with a history of palliative systemic chemotherapy were excluded from the study. The median age of the participants at diagnosis was 59.5 years, ranging from 53 to 76 years. Among the cohort, four patients were identified with hepatitis B infection, and one individual presented with co-infection of hepatitis B and C. Three participants were categorized as having non-B, non-C hepatitis, caused by alcoholic liver disease in one case and hepatic steatosis in two cases (Supplementary Table S1). Regarding radiotherapy approaches, five patients (63%) were treated with 3-dimensional conformal radiotherapy (3DCRT) at doses between 39-45 Gy in 10-18 fractions. Two patients (25%) received intensity-modulated radiotherapy (IMRT) at dose of 50 Gy in 25 fractions, and one patient (13%) underwent SBRT at dose of 50 Gy in 5 fractions (Supplementary Table S1). Over the course of the study, four patients succumbed to their illness. Tissue samples and relevant clinical data were collected with informed consent from all participants. The Institutional Review Board of Kangnam Sacred Heart Hospital (approval number HKS 2022-12-015, Seoul, South Korea) approved the study protocol, ensuring compliance with the ethical principles outlined in the Declaration of Helsinki.

### Generation of GPC3, luciferase expressing and GPC3 knockout stable cell line

The pLenti-C-mGFP-PP2A-Puro plasmid containing the full-length human GPC3 gene and the pLenti-C-Luciferase-P2A-Puro plasmid were purchased from OriGene (Rockville, MD, USA). Virus particles were generated following established protocols previously described (25). SNU449 cells were transduced using the Lenti-X accelerator starter kit (TaKaRa Bio Inc., Shiga, Japan) according to the manufacturer’s instructions. GPC3 and luciferase-expressing clones were obtained after 7-10 days of puromycin (2 µg/mL; Thermo Fisher Scientific) selection. Expression of GPC3 or luciferase was confirmed by Western blotting, flow cytometry, and immunofluorescence staining in each clone. Two single-cell clones were selected for further experiments: S4GA3 for SNU449/GPC3 and SV3 for SNU449/Vector (Supplementary Fig. S8). Additionally, a GPC3 knockout (GPC3^-^) Huh7 cell line was generated using a previously established method (Supplementary Fig. S9) (25).

### Quantitative PCR

Total RNA extraction and cDNA preparation were performed using the RNeasy Plus mini (Qiagen, Valencia, CA, USA) and QuantiTect reverse transcription (Qiagen) kits. To evaluate the level of GPC3 expression, quantitative PCR (qPCR) was performed on an ABI 7500 system (Applied Biosystems, Foster City, CA, USA) using a predesigned primer/probe set (Hs01018936_m1, Thermo Fisher Scientific) with the TaqMan® Universal PCR Master Mix (Applied Biosystems). following the manufacturer’s recommendations. The change of *GPC3* mRNA expression was normalized to endogenous ACTB. Negative and positive controls were included for the target gene in each assay.

### Western blotting

Western blotting was performed as previously described (25). The transferred proteins on the membrane were incubated with specific primary antibodies, including GPC3 (1:1000, 261M-96, Cell Marque, Rocklin, CA, USA), mGFP (1:1000, TA150122, OriGene), luciferase (1:1000, sc-74548, Santa Cruz Biotechnology), or glyceraldehyde-3-phosphate dehydrogenase (GAPDH, 1:5000, #CB1001, Sigma-Aldrich). Additional antibodies for ataxia-telangiectasia mutated (ATM, 1:1000, #2873), phospho-ATM (1:1000, #13050), checkpoint kinase 2 (CHK2, 1:1000, #3440), phospho-CHK2 (1:1000, #2197), cell division cycle 25C (CDC25C, 1:1000, #4688), phospho-CDC25C (1:1000, #4901), cell division cycle 2 (CDC2, 1:1000, #77055, aka, cyclin-dependant kinase 1, CDK1), phospho-CDC2 (1:1000, #4539), and p21 (1:1000, #2947) were purchased from Cell Signaling Technology (Danvers, MA, USA) and used for Western blotting, followed by appropriate secondary antibodies conjugated to horseradish peroxidase. Some membranes were stripped using Restore Plus Western Blot Stripping Buffer (Thermo Fisher Scientific) and reprobed. Immunoreactive bands were visualized by incubating the membranes with Clarity or Clarity Max ECL substrate (Bio-Rad, Hercules, CA, USA) and exposing them in a ChemiDoc MP imaging system (Bio-Rad). The intensity of the Western blot signals was quantified using ImageJ software (NIH, Bethesda, MD, USA) and the relative expression of each molecule was normalized to GAPDH.

### Flow cytometric analysis

To assess the expression of membrane-bound GPC3, cells from each culture were harvested using trypsin-EDTA, labeled with an APC-conjugated anti-GPC3 antibody (clone 1G12; Cell Marque) for 1 h on ice and then washed twice with an ice-cold staining buffer (1% (w/v) BSA and 0.5 mM EDTA in PBS). The stained cells were analyzed using a CytoFLEX Flow Cytometer (Beckman Coulter, Brea, CA, USA) within 30 minutes.

To evaluate cell death mechanisms, including necrosis and apoptosis, cells were stained with Annexin V-APC and propidium iodide (PI) according to the manufacturer’s instructions (BioLegend, San Diego, CA, USA). Additionally, cell cycle analysis was performed by measuring the DNA content. Cells were collected at 24, 48, and 72 h after irradiation, fixed in 70% ethanol at 4°C overnight, washed twice with the staining buffer, and then resuspended in a solution containing 100 μg/mL RNase A and 50 μg/mL PI (BioLegend). The flow cytometric data were acquired and processed using FlowJo software (FlowJo LLC, Ashland, OR, USA).

### Immunofluorescence

Approximately 5×10³ cells were plated and grown on Labtek II chamber slides (Nalgene Nunc, Naperville, IL, USA). The chamber slides were pre-rinsed with PBS, fixed using 4% paraformaldehyde (Sigma-Aldrich) for 10 minutes and permeabilized with 0.5% Triton X-100 (Sigma-Aldrich) for 1 minute, and subsequently blocked with 5% normal goat serum (Vector Laboratories, Burlingame, CA, USA) for 30 minutes at room temperature. To confirm membranous GPC3 expression, cells were labeled with a mouse anti-GPC3 antibody (clone 1G12; Cell Marque, 1:200 dilution in 2% BSA in PBS) at 4°C for overnight. For DNA damage assessment, cells were incubated with a mouse monoclonal anti-γH2AX antibody (Ser139, clone JBW301; Sigma-Aldrich, 1:400 dilution) under the same conditions. The following day, secondary antibodies conjugated to Alexa Fluor 555 (A-21422, Thermo Fisher Scientific, 1:500 dilution) were applied for 2 h at room temperature. The cells were counterstained with DAPI (1:1000, Thermo Fisher Scientific) for 30 minutes, washed three times, and mounted using ProLong Diamond Antifade (Thermo Fisher Scientific). and the cells were visualized using a Zeiss Axio Imager 2 microscope (Carl Zeiss Microscopy, Jena, Germany). The number of γH2AX foci per cell was manually scored in more than 50 cells per condition.

### IncuCyte® cell proliferation assay

Cell growth curves for the indicated cell lines were generated using the IncuCyte® SX5 live-cell imaging system (Sartorius, Göttingen, Germany). Approximately 3,000 cells were seeded in quadruplicate into 24-well plates. After an initial 24 h incubation, the cells were exposed to 6 Gy of radiation. Cell growth was subsequently monitored in a humidified atmosphere set at 37 °C with 5% CO_2_. Observations began 24 h post-IR and continued over the specified time period.

### Clonogenic survival assay

Radiosensitivity was evaluated by the clonogenic survival assay. Single-cell suspensions from cell culture were prepared by trypsinization. Designated numbers of cells were plated into six-well tissue culture plates and allowed to attach for 16 h. Following cell adhesion, the cells were irradiated at specific doses (204.5 cGy/min) using an X-Rad 320 irradiator (Precision X-ray, Inc., Branford, CT, USA). Radiation doses ranged between 0 and 8 Gy. After 12 to 14 days of incubation, the plates were stained with 0.5% crystal violet, and colonies containing at least 50 cells were counted. Surviving fractions were calculated based on the colony counts, with results presented as the mean ± SD from at least three independent experiments. The DMF was calculated from survival curves by determining the ratio of radiation doses required for 10% survival (GPC3^+^ dose divided by GPC3^-^ dose). A DMF value greater than 1 indicates a radioprotective effect.

### DNA strand break detection by alkaline comet assay

The alkaline comet assay was conducted following the manufacturer’s instructions (Trevigen, Gaithersburg, MD, USA). Cells were combined with molten agarose and layered onto CometSlides (Trevigen, Gaithersburg, MD, USA). The slides were incubated in a chilled lysis solution (Trevigen) for 1 hr and then treated with an alkaline unwinding solution (0.2 M NaOH and 1 mM EDTA) for 20 minutes. Alkaline electrophoresis was performed using a buffer containing 0.2 M NaOH and 500 mM EDTA (pH 8.2). After electrophoresis, slides were rinsed with distilled water, dehydrated in 70% ethanol, and allowed to dry overnight. DNA was stained with 100 µL of 1X SYBR Gold (Trevigen) for 30 minutes. Following staining, the slides were washed, air-dried, and imaged using a Zeiss Axio Imager 2 microscope. Quantification of comet tail DNA was performed using CometScore 2.0 software, applying the auto-threshold function to identify low-frequency DNA damage.

### RNASeq data analysis

To identify the effects of GPC3 on radioresistance in liver cancer cells, we performed RNA sequencing in GPC3^+^ and GPC3^-^ HepG2 and GPC3^+^ and GPC3^-^ Hep3B cell lines and analyzed gene expression. RNA samples from the engineered cell lines, collected 6 h after exposure to IR (6 Gy), were pooled and sequenced on the NovaSeq 6000 S1 platform using the Illumina Stranded Total RNA Prep and RiboZero Plus ligation methods, with paired-end sequencing in the NIH CCR-sequencing facility. The raw sequencing data underwent standard processing steps, including demultiplexing using Bcl2fastq (v2.20), adapter and quality trimming using Cutadapt, genome alignment to hg38 using the STAR aligner, and gene/isoform quantification with RSEM (v1.3.1) and Picard (v2.18.26), as described previously (25). Differential gene expression analysis was carried out using edgeR (v3.40.0), and genes with a false-discovery corrected p-value (p_adj_) < 0.05 and an absolute fold change > 1.0 were considered differentially expressed. The R package ComplexHeatmap (v.2.14.0) was used to generate gene expression heatmaps, while the R package clusterProfiler (v4.6.2) and GOrilla (55) were utilized for gene ontology, molecular function, and cell composition enrichment analysis, as well as to generate gene set enrichment analysis (GSEA) plots. The normalized gene counts derived from the STAR alignment were used for the GSEA analysis, with the log_2_ ratio as the enrichment metric. Additionally, differentially expressed genes (DEGs) from transcriptome profiling were analyzed using Ingenuity Pathway Analysis (IPA; QIAGEN). The core pathway analysis tool within IPA was employed to investigate gene ontology features and biological networks following radiation treatment, applying criteria of fold change ≥ ± 1.5 and *p*-value <0.05.

### Xenograft tumor models

All animal experiments were conducted following established ethical standards under a protocol approved by the Institutional Animal Care and Use Committee (protocol ROB-104, National Institutes of Health) and adhered to the guidelines set by the Institute of Laboratory Animal Resources, National Research Council. Female athymic nu/nu mice, aged 8 to 10 weeks, were obtained from Charles River Laboratories (Wilmington, MA, USA). A total of 5×10⁶ GPC3⁺ HepG2/Luc, GPC3^+^ A431/Luc and GPC3^-^ A431/Luc cells were subcutaneously injected into the right thigh of each mouse. For the GPC3⁻ HepG2/Luc group, 7×10⁶ cells were injected to generate tumors of comparable size to those formed by GPC3⁺ HepG2/Luc cells at the time of irradiation. For IR, mice were placed in a custom-designed lucite restrainer with lead shielding, ensuring targeted tumor exposure. A radiation dose of 10 Gy was administered to the right hind leg using an X-RAD 320 irradiator (Precision X-Ray) with 2.0 mm aluminum filtration at 320 kVp, delivered at a rate of 3.35 Gy/min. Tumor volumes were monitored as previously outlined (25), and bioluminescence (BL) imaging was performed 2-3 times weekly during the study period. For BL imaging, mice were anesthetized using 2% isoflurane, and images were captured 10 minutes after intraperitoneal injection of D-luciferin (150 mg/kg; Revvity) with the IVIS® Lumina Series III imaging system (Perkin Elmer, Waltham, MA, USA). Luminescence intensity within the defined tumor region of interest (ROI) was quantified using Living Image software (version 4.7.2, Perkin Elmer).

### Immunohistochemistry

Immunohistochemical analysis was performed on 5 µm sections of formalin-fixed, paraffin-embedded xenograft or human HCC tissues. The sections were deparaffinized using xylenes and rehydrated through a graded series of decreasing alcohol concentrations. Antigen retrieval was carried out with a pH 6.0 antigen retrieval buffer (Dako, Carpinteria, CA, USA). Endogenous peroxidase activity was blocked by incubating the sections with 3% hydrogen peroxide for 15 minutes. The xenograft tissue sections were then incubated for 1 hr at room temperature with either a rabbit monoclonal anti-GPC3 antibody (clone SP86; 1:3000 dilution; Abcam, Cambridge, MA, USA) or a rabbit monoclonal anti-Ki-67 antibody (clone SP6; 1:100 dilution; Abcam). Human tissue specimens were incubated with a mouse monoclonal anti-human GPC3 antibody (clone 1G12; Cell Marque) under the same conditions. Antigen-antibody reactions were visualized using the Dako Envision+ peroxidase system with 3,3-diaminobenzidine (DAB, Dako). Finally, the slides were digitized at 40× magnification using an Aperio AT2 digital scanner (Leica Biosystems, Wetzlar, Germany).

### Study Approval and Content Information

This study retrospectively analyzed tissue specimens from eight individuals diagnosed with hepatocellular carcinoma who received radiotherapy between September 2013 and December 2020 at Kangnam Sacred Heart Hospital. Tissue samples and associated clinical information were obtained with written informed consent from all participants. The study protocol was reviewed and approved by the Institutional Review Board of Kangnam Sacred Heart Hospital (HKS 2022-12-015) and conducted in accordance with the ethical principles of the Declaration of Helsinki. All animal experiments were performed following institutional and national guidelines. Protocols were approved by the National Institutes of Health Institutional Animal Care and Use Committee and adhered to the recommendations of the Institute of Laboratory Animal Resources, National Research Council.

### Statistical analysis

Data analysis was conducted using Prism 10.0 (GraphPad Software Inc., San Diego, CA, USA) and SPSS Statistics for Windows, version 29 (IBM Corp., Armonk, NY, USA). Results are presented as mean ± standard deviation (SD) or standard error of the mean (SEM), as indicated. Differences in proliferation, tumor volume, and bioluminescence at multiple time points were assessed using two-way ANOVA, with significant differences highlighted at the final time point in the plot. Comparisons between two groups were performed using two-tailed unpaired t-tests. Survival analysis employed OS model, treating all patient deaths as events and censoring other patients at their last recorded visit. The date of completion of radiotherapy was defined as time zero for survival time calculation. The Kaplan–Meier method was used to estimate survival curves, and group comparisons were made using the log-rank test. A *p*-value below 0.05 was considered statistically significant.

## Acknowledgments

This research was fully supported by the Intramural Research Program of the National Cancer Institute, National Institutes of Health (Molecular Imaging Branch, Center for Cancer Research). The contributions of the NIH authors were made as part of their official duties as NIH federal employees, are in compliance with agency policy requirements, and are considered Works of the United States Government. However, the findings and conclusions presented in this paper are those of the authors and do not necessarily reflect the views of the NIH or the U.S. Department of Health and Human Services. We also acknowledge the NIH Cyclotron Facility for the production and supply of the radioisotopes used in this study.

## Author contributions

JYC and FEE conceived and designed the experiments. JYC, HM, SK, JSK, DN, and ST conducted the *in vitro* experiments. OWL performed RNA-seq data analysis. WL, AOW, and EJC conducted the *in vivo* experiments. JYC, NA, SR, and SMH acquired and analyzed the IHC data. JYC, JWK, and FEE acquired and analyzed the clinical data. JYC drafted the manuscript. SMH, THS, DEC, PLC, and FEE reviewed and revised the manuscript. All authors contributed to and approved the final version of the manuscript.

## Data Availability Statement

The data that support the findings of this study are available from the Corresponding Authors, [Joon-Yong Chung and Freddy E. Escorcia], upon reasonable request.

