## Supplementary Figure S1-S9 and Table S1 for "Ablation of glypican-3 enhances radiosensitivity in liver cancer by prolonging G2/M arrest and activating the ATM/CHK2 pathway"

**Affiliations:**
[1] Molecular Imaging Branch, Center for Cancer Research, National Cancer Institute, National Institutes of Health, Bethesda, MD 20892, USA.
[2] Integrative Tumor Epidemiology Branch, Division of Cancer Epidemiology and Genetics, National Cancer Institute, National Institutes of Health, Bethesda, MD 20892, USA.
[3] Radiation Oncology Branch, Center for Cancer Research, National Cancer Institute, National Institutes of Health, Bethesda, MD 20892, USA.
[4] Department of Pathology, Kangnam Sacred Heart Hospital, Hallym University College of Medicine, Seoul, 07441, Republic of Korea
[5] Oxford Institute for Radiation Oncology, Department of Oncology, University of Oxford, Oxford, UK.
[6] Laboratory of Pathology, Center for Cancer Research, National Cancer Institute, National Institutes of Health, Bethesda, MD 20892, USA.

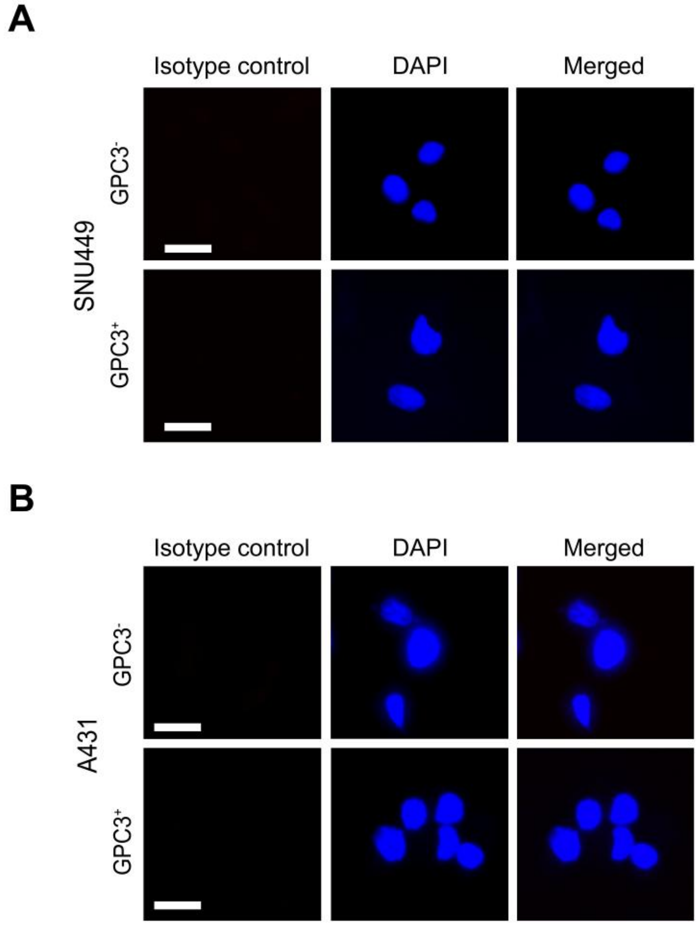


**Supplementary Fig. S1.** Immunofluorescence staining with isotype control antibody. The isotype control was negative in both SNU449/GPC3 (*A*) and A431/GPC3 (*B*) cells as well as in cells deficient in GPC3. Scale bars are shown for 50 $\mu$m.

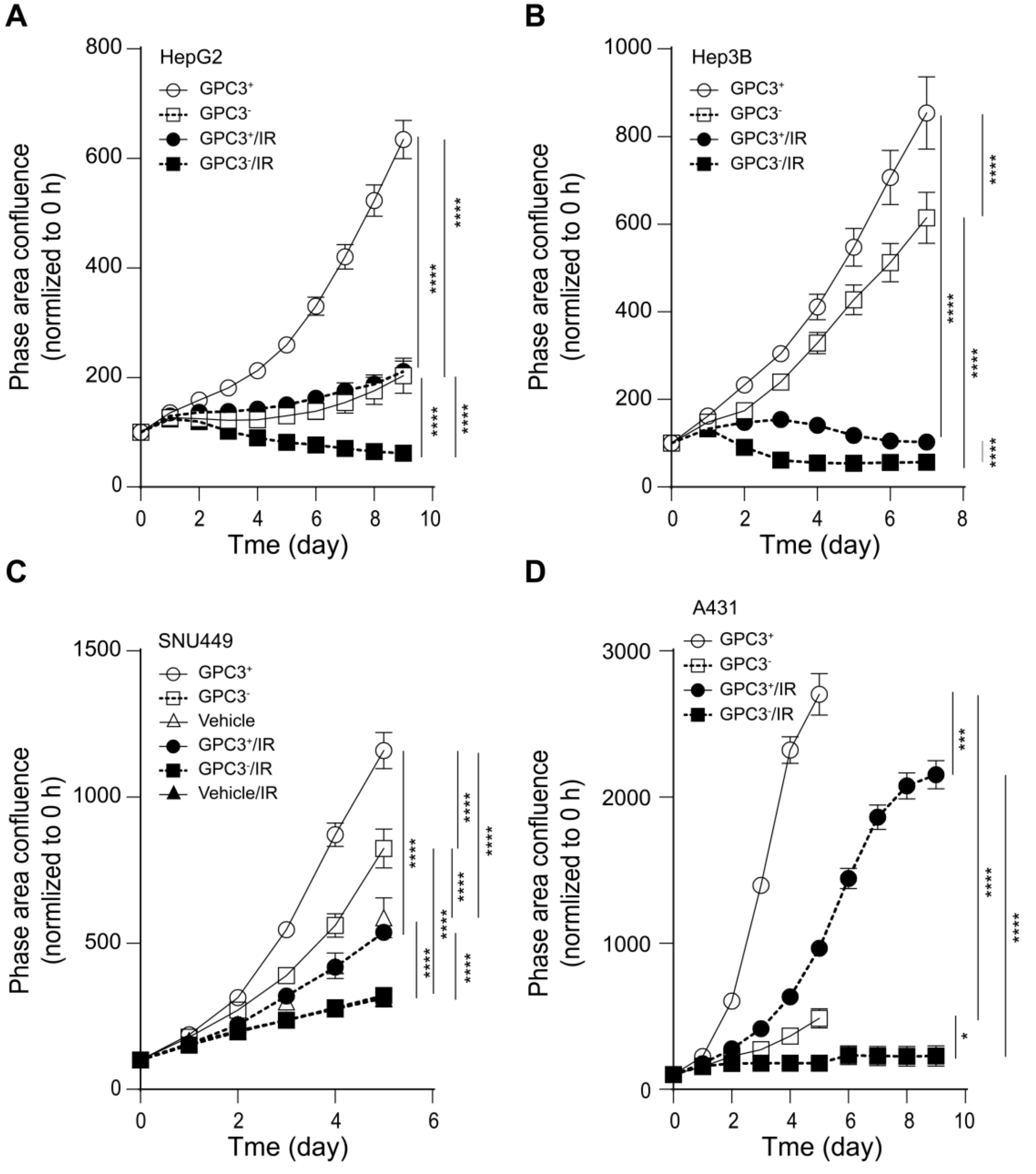


**Supplementary Fig. S2.** Proliferation analysis of GPC3$^+$ and GPC3$^-$ cancer cells under irradiated and non-irradiated conditions. Cell lines tested included HepG2 (*A*), Hep3B (*B*), SNU449 and vector control SNU449/V (*C*), and A431 (*D*), each with GPC3$^+$ and GPC3$^-$ variants. Cell growth was tracked using the IncuCyte® live-cell imaging system, both without treatment and after exposure to 6 Gy irradiation (IR). Statistical comparisons were made using two-way ANOVA followed by multiple comparisons. *$p$<0.05, ***$p$<0.001, ****$p$<0.0001.

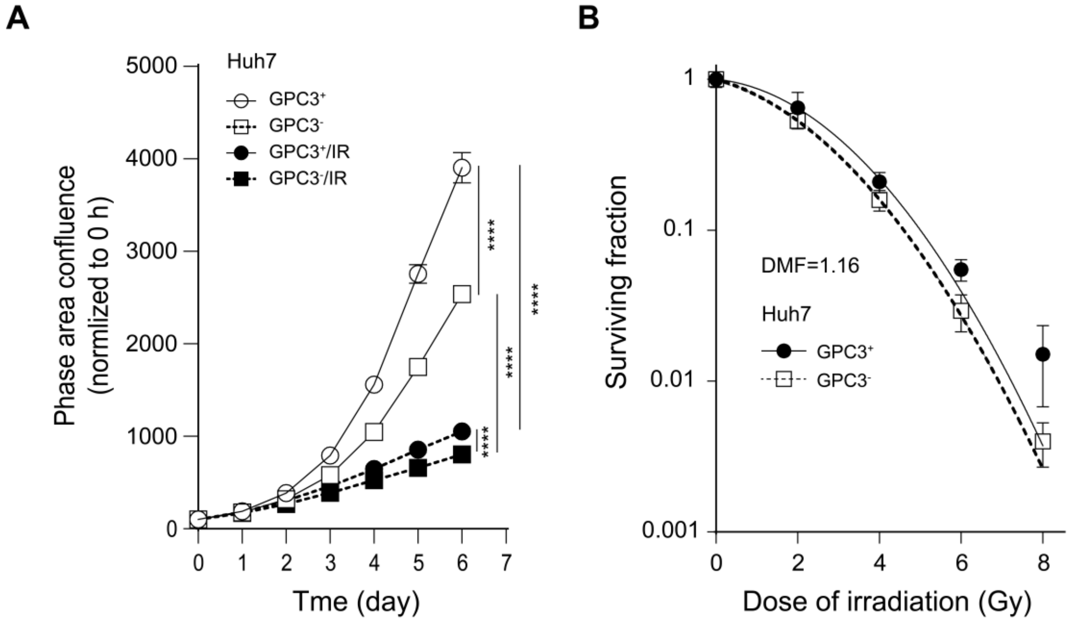


**Supplementary Fig. S3.** Analysis of proliferation and clonogenic capacity of $GPC3^{+}$ and $GPC3^{-}$ Huh7 cell lines. (*A*) Cell proliferation was monitored using the IncuCyte® live-cell imaging system under control conditions and after 6 Gy IR. (*B*) Clonogenic survival was assessed 10–14 days post-treatment, and survival curves were generated. GPC3-deficient cells showed a modest decrease in colony-forming ability, with a dose modification factor of 1.16 observed in Huh7 cells. Data represent mean survival fractions ± SD from three independent experiments.

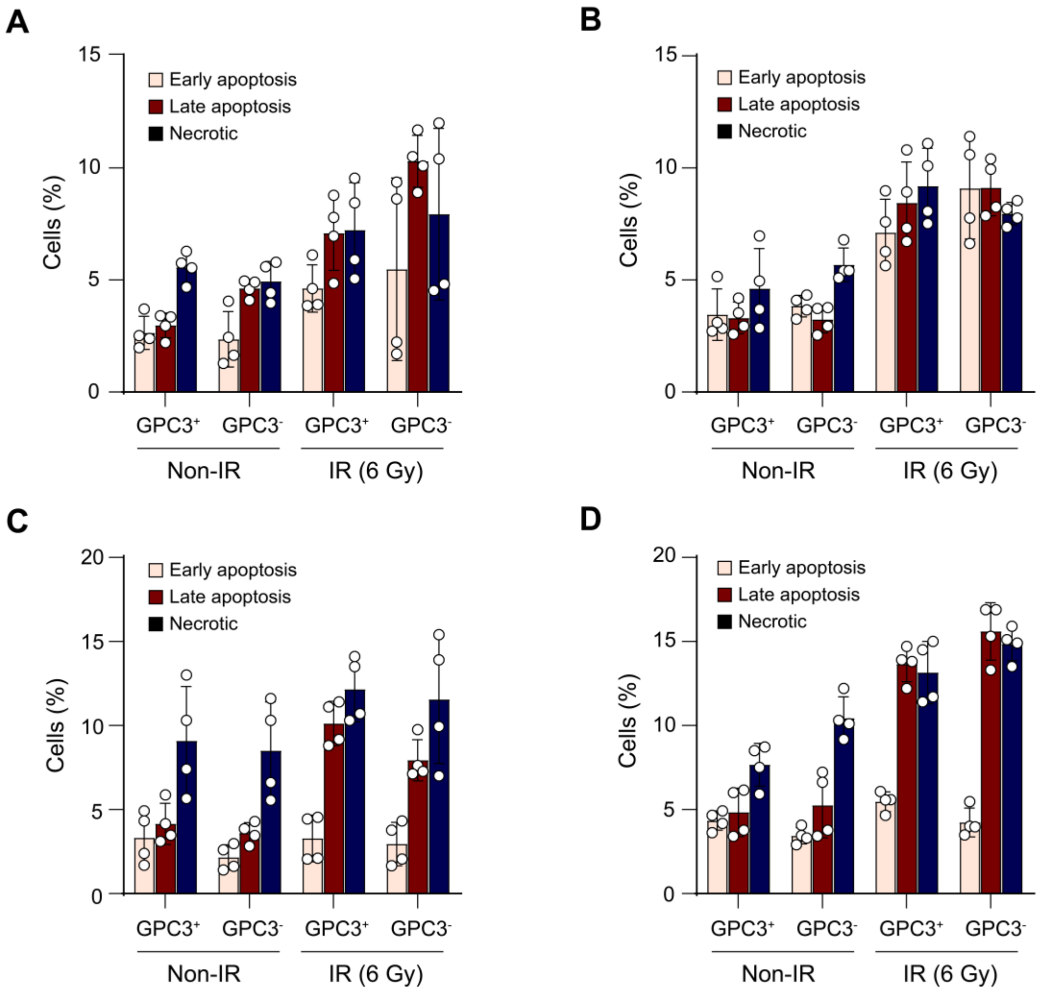


**Supplementary Fig. S4.** Apoptotic and necrotic profiling of GPC3⁺ and GPC3⁻ liver cancer cells following IR. Flow cytometric analysis was performed to assess apoptosis and necrosis in GPC3⁺ and GPC3⁻ HepG2 cells at 24 h (*A*) and 48 h (*B*) after exposure to 6 Gy of ionizing radiation. A similar analysis was conducted in GPC3⁺ and GPC3⁻ Hep3B cells at 24 h (*C*) and 48 h (*D*) post-irradiation.

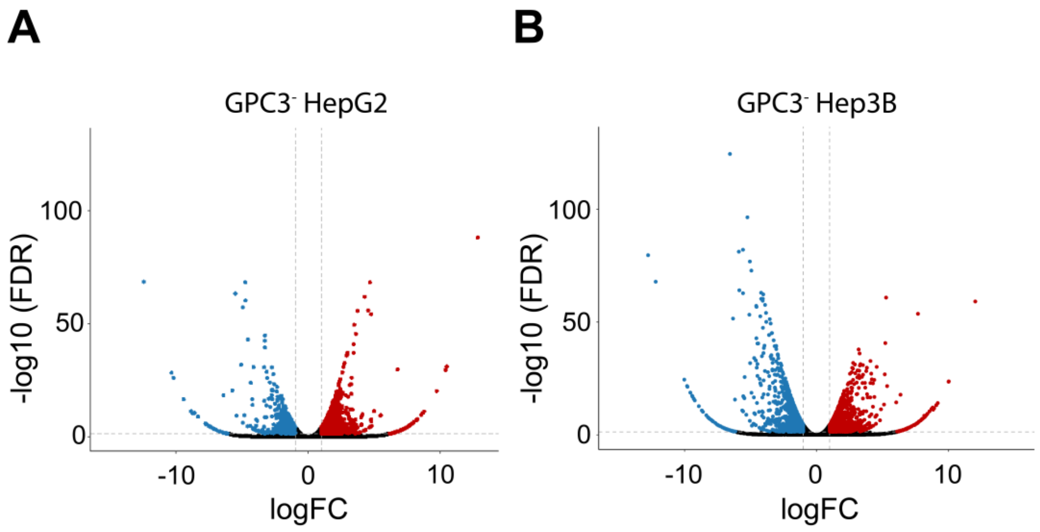


**Supplementary Fig. S5.** Differential gene expression in GPC3$^-$ liver cancer cell lines. Volcano plots illustrating the differential gene expression profiles in GPC3$^-$ HepG2 (*A*) and Hep3B (*B*) cells compared to their respective parental lines. Each plot displays log2 fold change (x-axis) versus -log10 FDR-adjusted *p*-value (y-axis) for individual genes. Genes with significant downregulation are highlighted in blue, while those with significant upregulation are highlighted in red.

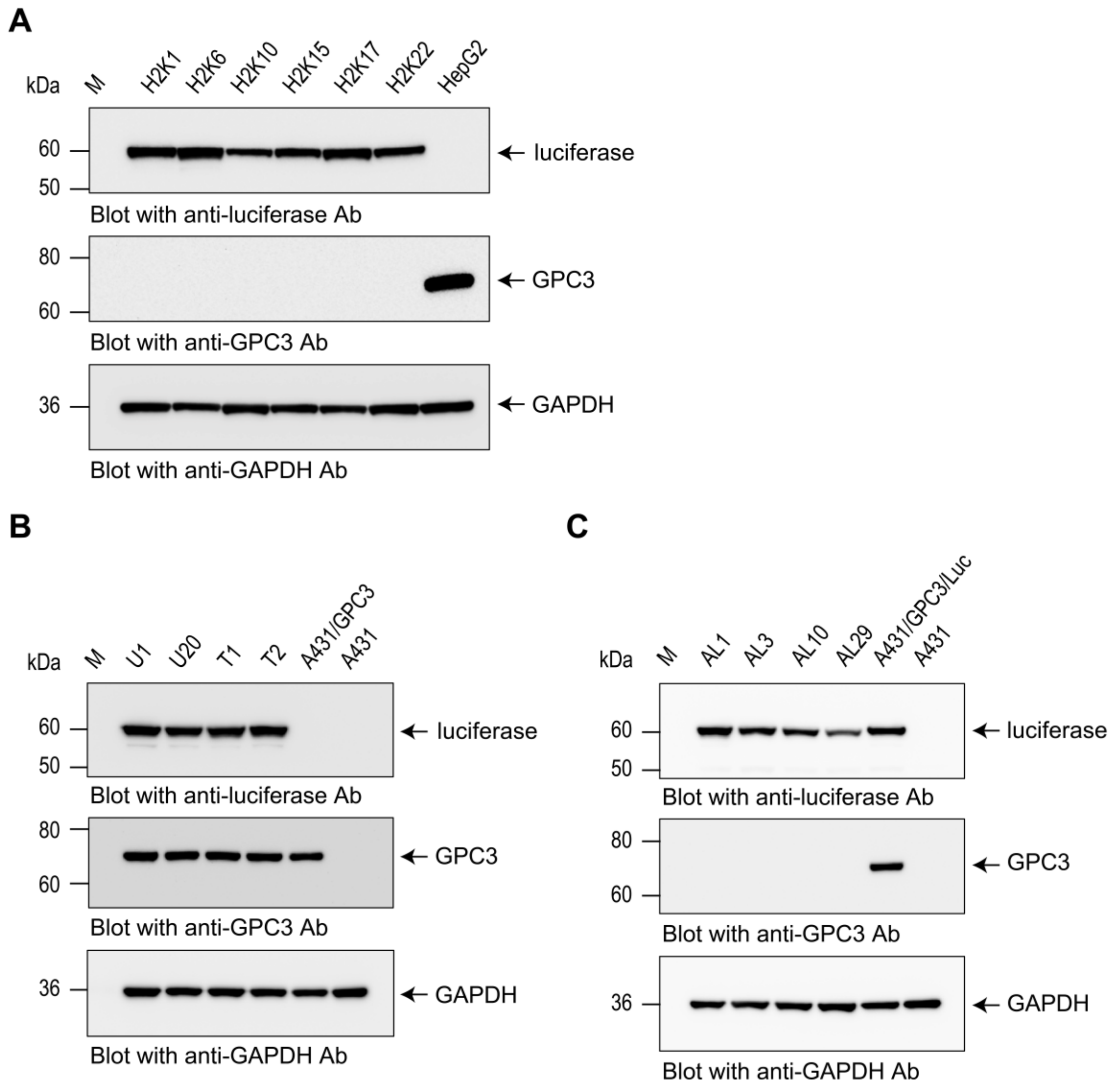


**Supplementary Fig. S6.** Overexpression of luciferase in GPC3$^-$ HepG2, A431/GPC3, and A431 cells using a lentiviral system. Single-cell clones H2K17 (*A*), T2 (*B*), and AL1 (*C*) were selected for GPC3$^-$ HepG2, A431/GPC3, and A431 cells, respectively. These clones were used for *in vivo* experiments. GAPDH was used as a loading control.

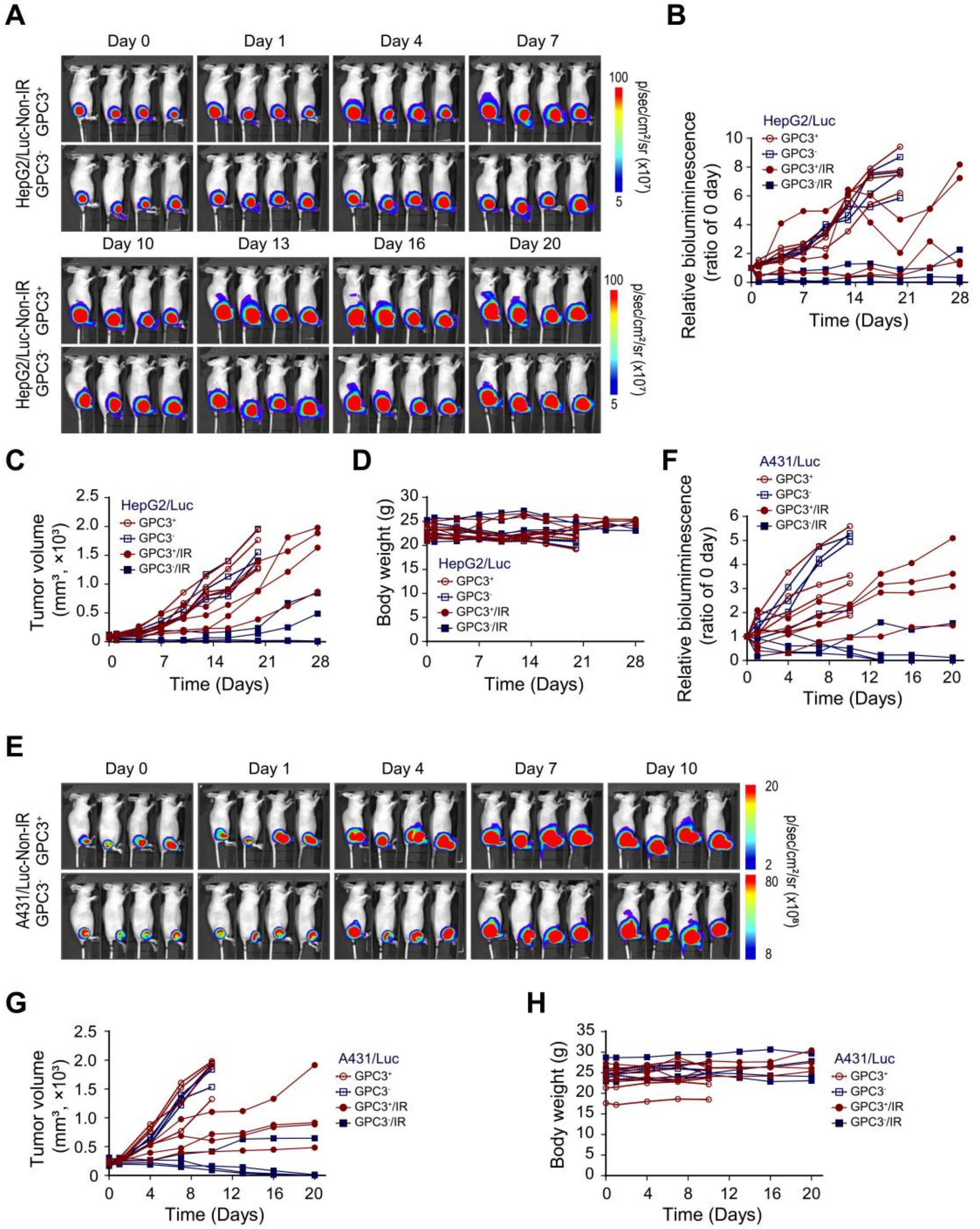


**Supplementary Fig. S7.** Tumor progression and body weight monitoring in mice bearing GPC3$^{+}$ and GPC3$^{-}$ tumors with and without irradiation (IR). (*A*, *E*) Longitudinal bioluminescence imaging of mice injected with GPC3$^{+}$ HepG2/Luc, GPC3$^{-}$ HepG2/Luc, GPC3$^{+}$ A431/Luc, or GPC3$^{-}$ A431/Luc cells, under non-IR conditions. (*B*, *F*) Quantification of bioluminescence signal over time in the same cohorts, under no-IR and IR conditions. (*C*, *G*) Tumor growth curves estimated by caliper measurements. (*D*, *H*) Body weight changes over time in mice injected with GPC3$^{+}$ HepG2/Luc, GPC3$^{-}$ HepG2/Luc, GPC3$^{+}$ A431/Luc, or GPC3$^{-}$ A431/Luc cells.

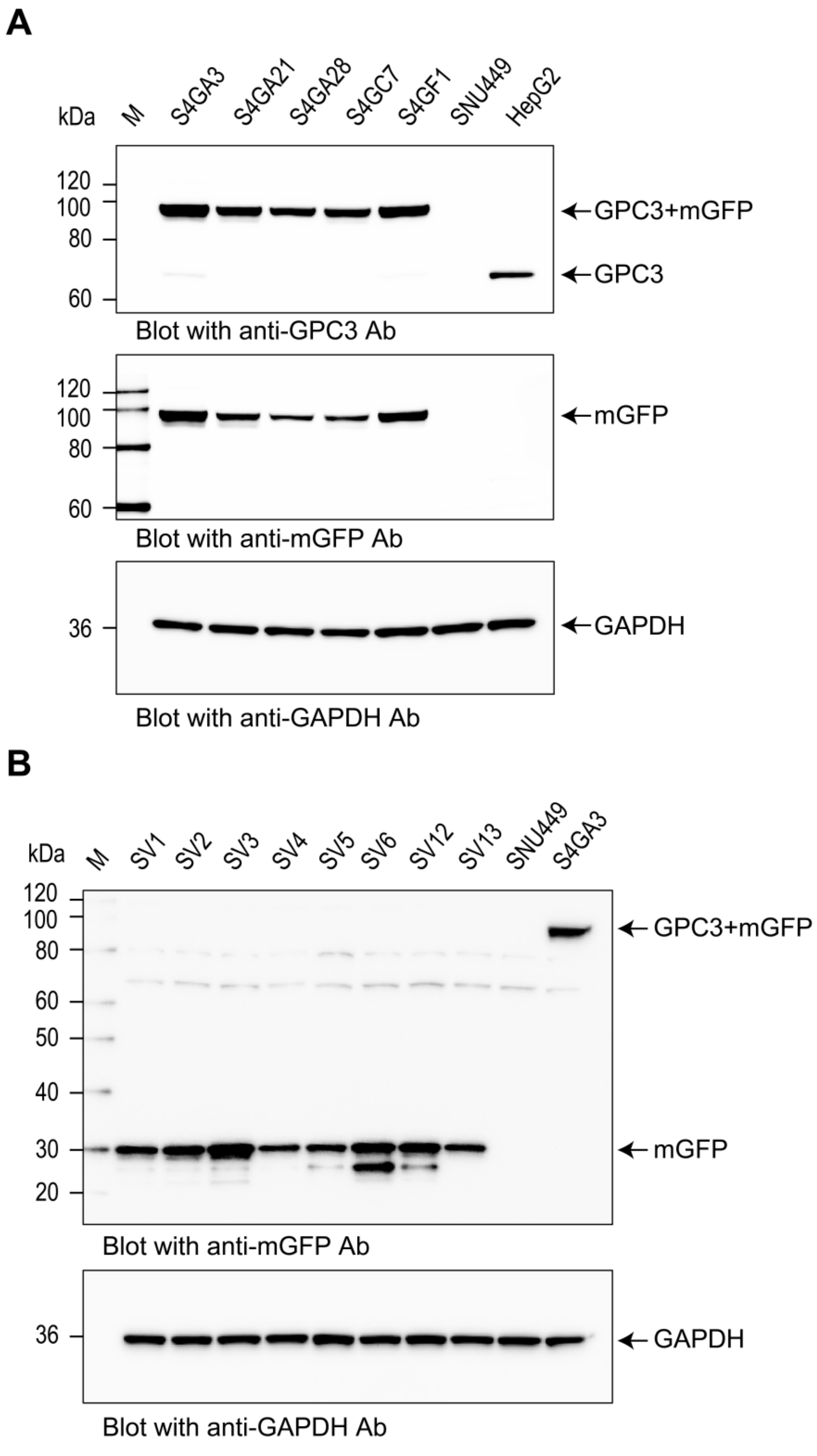


**Supplementary Fig. S8.** Overexpression of GPC3 in SNU449 cells using a lentiviral system. Single-cell clones S4GA3 (*A*) and SV3 (*B*) were selected for GPC3 overexpression and lentiviral vector control (vehicle), respectively. These clones were used for subsequent analyses. GAPDH was used as a loading control.

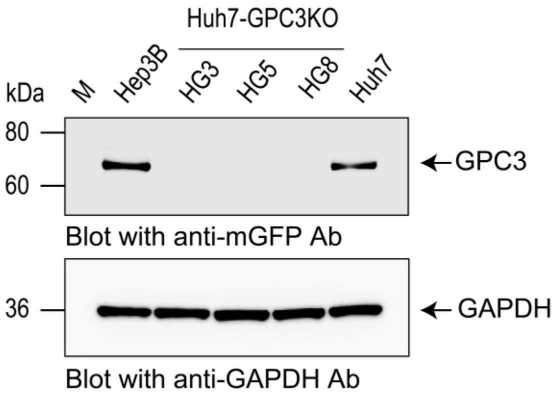


**Supplementary Fig. S9.** CRISPR/Cas9-mediated knockout of GPC3 in Huh7 cells. The HG3 clone was selected for further experiments.

**Supplementary Table S1.** Clinicopathological characteristics, GPC3 expression, and radiation therapy details of study cohort

| No. | Age | Sex | Growth type | T stage | N stage | Cirrhosis | Steatosis | HBV | HCV | GPC3 expression | RTx (total dose/fraction) |
|---|---|---|---|---|---|---|---|---|---|---|---|
| 1 | 72 | M | Expanding nodular | T3 | N0 | - | + | - | - | - | 3DCRT (30 Gy/10) |
| 2 | 62 | M | Nodular with perinodular expansion | T2 | N0 | + | - | + | + | - | 3DCRT (39 Gy/11) |
| 3 | 76 | M | Expanding nodular | T3 | N0 | - | + | - | - | - | SBRT (50Gy/5) |
| 4 | 53 | M | Nodular with perinodular expansion | T1 | N0 | + | + | + | - | + | 3DCRT (39 Gy/13) |
| 5 | 56 | M | Expanding nodular | T2 | N0 | - | - | + | - | - | 3DCRT (39 Gy/13) |
| 6 | 58 | M | Expanding nodular | T3 | N0 | + | + | + | - | + | IMRT (50 Gy/25) |
| 7 | 57 | M | Expanding nodular | T3 | N0 | + | - | + | - | + | 3DCRT (45 Gy/18) |
| 8 | 61 | M | Nodular with perinodular expansion | T2 | N0 | + | + | - | - | - | IMRT (50 Gy/25) |

HBV, hepatitis B virus; HCV, hepatitis C virus; 3DCRT, three-dimensional conformal radiation therapy; SBRT, stereotactic body radiation therapy; IMRT, intensity-modulated radiation therapy
